# High-speed volumetric single-molecule imaging using dual-wavelength light sheets and PSF-engineered enhanced biplane detection

**DOI:** 10.64898/2026.06.20.733419

**Authors:** Prakash Joshi, Nahima Saliba, Siyang Cheng, Yuya Nakatani, Dafei Xiao, Reut Orange-Kedem, Yoav Shechtman, Anna-Karin Gustavsson

## Abstract

Single-molecule localization microscopy (SMLM) enables nanoscale imaging but remains limited in three-dimensional (3D), high-speed, and high-density applications due to background fluorescence, photon inefficiency, and large point-spread function (PSF) footprints. Here, we present single-objective light-sheet microscopy with PSF-engineering enhanced biplane detection (SoLiD-3D), a versatile imaging platform that integrates dual-wavelength light-sheet illumination with dual-color, multi-configuration biplane imaging for parallel acquisition with PSF engineered detection for high-speed volumetric SMLM. Parallelized single-objective light-sheet excitation combined with PSF engineering overcomes key limitations of conventional wide-field and biplane approaches. Independent control of two excitation wavelengths for optical sectioning enables simultaneous dual-target imaging and single-target dual-color imaging with improved contrast and temporal resolution utilizing dynamically displaced light sheets for volumetric coverage. Using SoLiD-3D, we demonstrate high-speed single- and dual-target dual-color imaging that doubles localization density without sacrificing photon efficiency and continuous volumetric imaging via PSF-engineering enhanced biplane detection for whole-cell 3D imaging with improved axial localization performance over extended depth ranges. We further demonstrate improved speed by utilizing the Hummus PSF, a compact engineered PSF that enables high-precision 3D localization with a substantially reduced spatial footprint, for the first time for super-resolution imaging applications. Taken together, SoLiD-3D mitigates the trade-off between axial range and localization precision and offers improved speed compared to conventional 3D SMLM approaches.

## Introduction

Three-dimensional (3D) fluorescence microscopy has become an essential tool for studying the spatial organization and dynamics of biological systems at cellular and subcellular levels. Single-molecule localization microscopy (SMLM)^1^ and other super-resolution imaging techniques^2–5^ have enabled imaging beyond the diffraction limit, revealing nanoscale structural and functional details that are otherwise unresolvable with conventional optical microscopy approaches. Despite these advances, extending super-resolution microscopy to volumetric imaging remains challenging due to inherent trade-offs between axial resolution, imaging speed, photobleaching, and background fluorescence. Light-sheet (LS) fluorescence microscopy has emerged as a powerful strategy to address many of these limitations^2–4,6–9^. By confining excitation to a thin plane within the sample, LS illumination provides intrinsic optical sectioning which reduces out-of-focus excitation and photobleaching, and improves the signal-to-background ratio (SBR), making it highly effective for 3D biological imaging^10–13^. Numerous implementations of LS microscopy have been developed, including selective plane illumination microscopy (SPIM)^14,15^, oblique plane microscopy (OPM)^16^, highly inclined and laminated optical sheet (HILO)^17^ microscopy, single-objective LS microscopy^18^, and scanned Gaussian- or Bessel-beam–based approaches^19,20^, each offering distinct trade-offs between resolution, field of view, and imaging speed. Several approaches have been developed to enable volumetric SMLM, including interferometric detection^21^, multi-objective geometries^22–25^, and engineered point-spread functions (PSFs) generated via phase modulation in the Fourier plane^10,12,21,26–30^. Among these, engineered PSFs, such as the double-helix (DH) or tetrapod PSFs, offer a flexible and hardware-efficient means of encoding axial position over extended depth ranges while remaining compatible with high-NA single-objective detection architectures^31–34^. However, when combined with conventional wide-field epi-illumination, localization of engineered PSFs with extended axial ranges is often difficult due to increased background fluorescence and photobleaching, which degrades localization precision and spatial or temporal resolution. Biplane detection has been introduced as a complementary strategy for volumetric imaging, enabling simultaneous acquisition of two axially separated focal planes^29^. This approach improves axial sampling and temporal resolution and can be implemented using relatively simple optical modifications^27,35^. However, conventional biplane systems typically rely on splitting the emission signal into two channels, which can compromise localization precision. What is needed for improved whole-cell SMLM is a platform that offers fast, photon-efficient, and low-background 3D imaging for improved precision and speed.

In this work, we present single-objective LS microscopy with PSF-engineering enhanced biplane detection (SoLiD-3D), a versatile imaging platform that integrates dual-wavelength LS illumination and multi-configuration dual-color biplane imaging with engineered-PSF detection for high-speed volumetric SMLM. By combining LS excitation with parallelized PSF engineering and biplane detection strategies, our system overcomes key limitations associated with conventional widefield epi-illumination and emission collection in SMLM. The illumination pathway provides independent control of two spectrally distinct LSs, enabling simultaneous dual-target imaging and single-target dual-color imaging using tunable offsets between LS planes for extended volumetric coverage and increased speed without sacrificing SBR. The detection pathway employs a high-NA objective lens for efficient photon collection and PSF engineering in two independent color-channels for parallel 3D localization, where the detection planes can be flexibly offset by translation of a single lens to enable biplane detection, thus extending the axial range while benefitting from the high performance of short-axial range PSFs in each color channel. Furthermore, by splitting the emission photons spectrally instead of non-selectively, SoLiD-3D offers this PSF-engineering enhanced biplane detection without compromising photon detection efficiency. Such optical modifications enable seamless transition between multicolor, multi-plane, and engineered-PSF imaging modalities utilizing a single camera.

We show that the background suppression when using our LSs improves the 3D localization precision for cellular imaging compared with conventional epi-illumination. Next, we demonstrate that SoLiD-3D improves the precision and throughput of single-plane single-target imaging by enabling two-color LS illumination and parallel detection of the same structure in two color channels, and of whole-cell single-target imaging by enabling tunable LS offsets and biplane detection of two spatially offset planes with the retained performance of short axial range engineered PSF in each plane. We also show that SoLiD-3D enables parallelized and high-performance dual-target imaging.

To further improve the speed and performance in dense imaging conditions, we leverage the Hummus PSF^36,37^, which is a compact engineered PSF that enables high-precision 3D localization with a substantially reduced lateral footprint. Unlike widely used engineered PSFs such as the DH-PSF^33^ or Tetrapod PSFs^38,34^, which suffer from reduced performance at high emitter densities^10^, the Hummus PSF^36,37^, optimally designed for high density SMLM, confines the emitter signal into a compact region while retaining axial sensitivity, enabling imaging at higher emitter concentrations. Here, we demonstrate the Hummus PSF for the first time for super-resolution imaging with over 2x speed compared with a DH-PSF with the same axial range^33^ while preserving high spatial resolution, illustrating that its reduced footprint enables robust 3D localization in dense, biologically relevant environments where spatially extended PSFs are prone to overlap and localization failure.

Taken together, SoLiD-3D provides a versatile and powerful tool for studying complex cellular architectures and molecular interactions at the nanoscale with improved precision and speed.

## Results

### SoLiD-3D design

SoLiD-3D, a platform that integrates single-objective LS excitation with spectrally separated 3D detection enables whole-cell imaging with high spatiotemporal resolution (Fig. 1a, Supplementary Fig. 1). SoLiD-3D employs two independently controlled LSs in the excitation path, which can be flexibly positioned at distinct image planes. This configuration effectively reduces fluorescence background for each wavelength and each selected plane, improving the SBR for high-fidelity volumetric imaging. Rather than dividing photons with a beam splitter, SoLiD-3D separates detection channels spectrally, thereby preserving the total photon counts in each channel (Fig. 1b, c).

**Figure 1.**
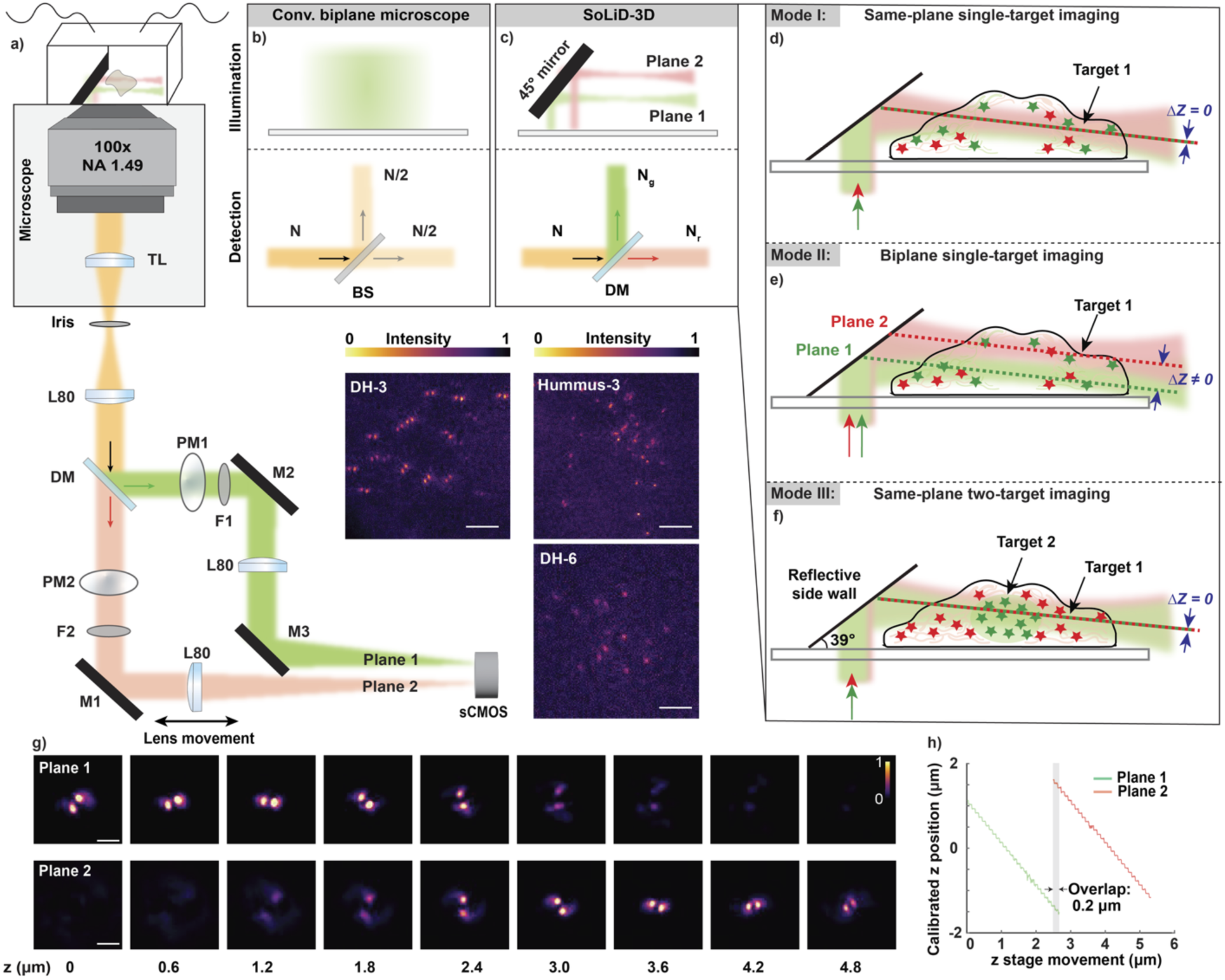
Single-objective light-sheet microscopy with PSF-engineering enhanced biplane detection (SoLiD-3D). (a) Simplified schematic of the SoLiD-3D microscopy platform integrating two-color single-objective light-sheet (LS) illumination and biplane detection with engineered point-spread functions (PSFs) in each detection path. Scale bars: 5 µm. (b,c) Two excitation and detection planes (Plane 1 and Plane 2), which can be co-aligned or axially offset, are spectrally separated using a dichroic mirror, eliminating the 50/50 photon split inherent to beam-splitter based biplane implementations. The spectral separation enables imaging of (d) the same focal plane for high-density localization, (e) two focal planes for parallelized whole-cell imaging with improved speed, or (f) simultaneous two-target imaging within a single acquisition. The schematics are not to scale. (g) Experimental double-helix (DH) PSFs acquired by imaging of fluorescent beads using ∼2.7 µm axial-range DH-PSFs (DH-3). Scale bars: 2 µm. (h) The two planes are here offset to have an overlap of ∼200 nm. This allows simultaneous detection of the full cellular thickness (∼5 µm), while ensuring continuous axial sampling and registration between the two planes. Abbreviations: BS: beam splitter, DM: dichroic mirror, F: filter, L: lens, M: mirror, N: total number of photons, N_g_: photons in the green channel, N_r_: photons in the red channel, PM: phase mask, TL: tube lens.

Two reflected, steerable, and dithered single-objective LSs of 560 nm and 647 nm were generated as described previously^10,20^, with thicknesses of 1.2 μm (1/e^2^ beam waist radius) (Supplementary Fig. 2), lateral widths of 56 μm and 48 μm (1/e^2^ diameter) (Supplementary Fig. 3), and confocal parameters of 15 μm and 13 μm (Supplementary Fig. 4), respectively, ensuring selective excitation within the axial range of the used PSFs with suitable parameters for cell imaging. To spatially align the excitation volumes of both LSs, galvanometric mirrors and an electrically tunable lens was incorporated into each illumination path, enabling precise translation of the beam waist. By varying the current applied to the tunable lens from 0 mA to −120 mA, the focal positions of both LSs could be synchronously shifted over a range of ∼0–50 μm, allowing the confocal parameters of the two LSs to overlap at the same imaging plane throughout the imaging volume (Supplementary Fig. 5).

The same high-NA used to generate the LSs was also used to collect the light emitted from the sample. The collected light was then spectrally separated into two different paths, “green” and “red”, each containing a 4f relay system for PSF engineering. The second 4f lens in the red path can be translated along the optical axis using a translational stage to enable tunable offsets between the detection planes of the two channels (Fig. 1a), facilitating PSF-engineering enhanced biplane detection.

The system incorporates a micro-mirror incorporated microfluidic system for LS reflection into the sample and automated solution exchange and solution perfusion, enabling dynamic experiments and multiplexed imaging strategies without manual intervention^10,39–41^.

SoLiD-3D’s highly versatile platform is demonstrated here for three imaging modalities:

i. single-plane dual-color single-target imaging, (ii) biplane dual-color single-target imaging, and
ii. single- or biplane dual-target imaging (Fig. 1d-f).

First, in the single-plane dual-color single-target modality, a single molecular target is illuminated by the two co-aligned LSs and detected simultaneously in two spectrally separated channels. This approach effectively doubles the number of detected emitters per frame without reducing photon counts per localization or SBR, thereby increasing localization density (Fig. 1d), and by extension, temporal resolution.

Secondly, in the biplane dual-color single-target imaging, one of the LSs (647 nm) and the second 4f lens in the red detection path is displaced, allowing simultaneous illumination and detection of two different planes (Fig. 1e). With this approach, the imaging depth is extended to cover the entire cell volume simultaneously, while retaining thin optical section for improved SBR and the axial precision of short axial range engineered PSF in each plane for further improved precision. Using DH-PSF with axial ranges of ∼ 2.7 µm (DH-3), the illumination and detection planes were offset to ensure an axial overlap of 200-300 nm (Fig. 1h) to allow for simultaneous whole-cell volumetric imaging (Supplementary Fig. 6).

Thirdly, in the dual-target modality, the two LSs are here spatially co-aligned and used to image two distinct molecular targets simultaneously throughout the same plane (Fig. 1f). This configuration increases the imaging speed 2-fold compared to sequential imaging while preserving photon collection efficiency and localization precision. This also enables high-contrast 3D imaging where simultaneous mapping of the two targets is critical, either in the same or in two different planes.

Taken together, this coordinated illumination-detection strategy eliminates unnecessary background excitation, reduces crosstalk, preserves photons, and enables efficient volumetric super-resolution imaging throughout whole cells.

### Single-target imaging with improved precision and speed

To enhance the imaging speed without compromising localization precision, we implemented co-aligned two-color LS illumination with simultaneous dual-color imaging with PSF engineering detection using DH phase masks in two co-aligned detection paths, each with an axial range of ∼2.7 µm (DH-3). We showcase the performance of SoLiD-3D for 3D SMLM by imaging the mitochondrial outer membrane protein TOMM20 labeled with Cy3B (green channel) and ATTO 647N (red channel) using DNA-PAINT imaging^40^, in which transient binding of fluorescently labeled imager strands generates stochastic single-molecule blinking events for localization microscopy. In this configuration, TOMM20 was detected in two spectrally separate channels concurrently, enabling parallelized dual-color acquisition of localization events. Using SoLiD-3D, the mitochondrial network was resolved with high structural fidelity in 3D in both color channels independently (Fig. 2). Upon merging the localizations from the two color channels, the resulting 3D reconstruction preserved the characteristic complex and branched morphology of TOMM20 across the imaged cell volume (Supplementary Videos 1 and 2). Analysis of 20,000 frames yielded ∼370,000, ∼310,000, and ∼680,000 localizations in the green, red, and merged channel, respectively, demonstrating an effective doubling of the localization density compared to single-channel acquisition. Because the two datasets were recorded in parallel rather than sequentially, the total acquisition time remained unchanged. Hence, this parallelization directly improves temporal and/or spatial resolution for volumetric imaging. The effective spatial resolution evaluated for the full merged reconstruction using Fourier Ring Correlation (FRC) analysis yielded a resulting resolution of 40/69/65 nm in the xy/yz/xz planes, respectively, from only 20,000 acquired frames (Supplementary Fig. 7).

**Figure 2.**
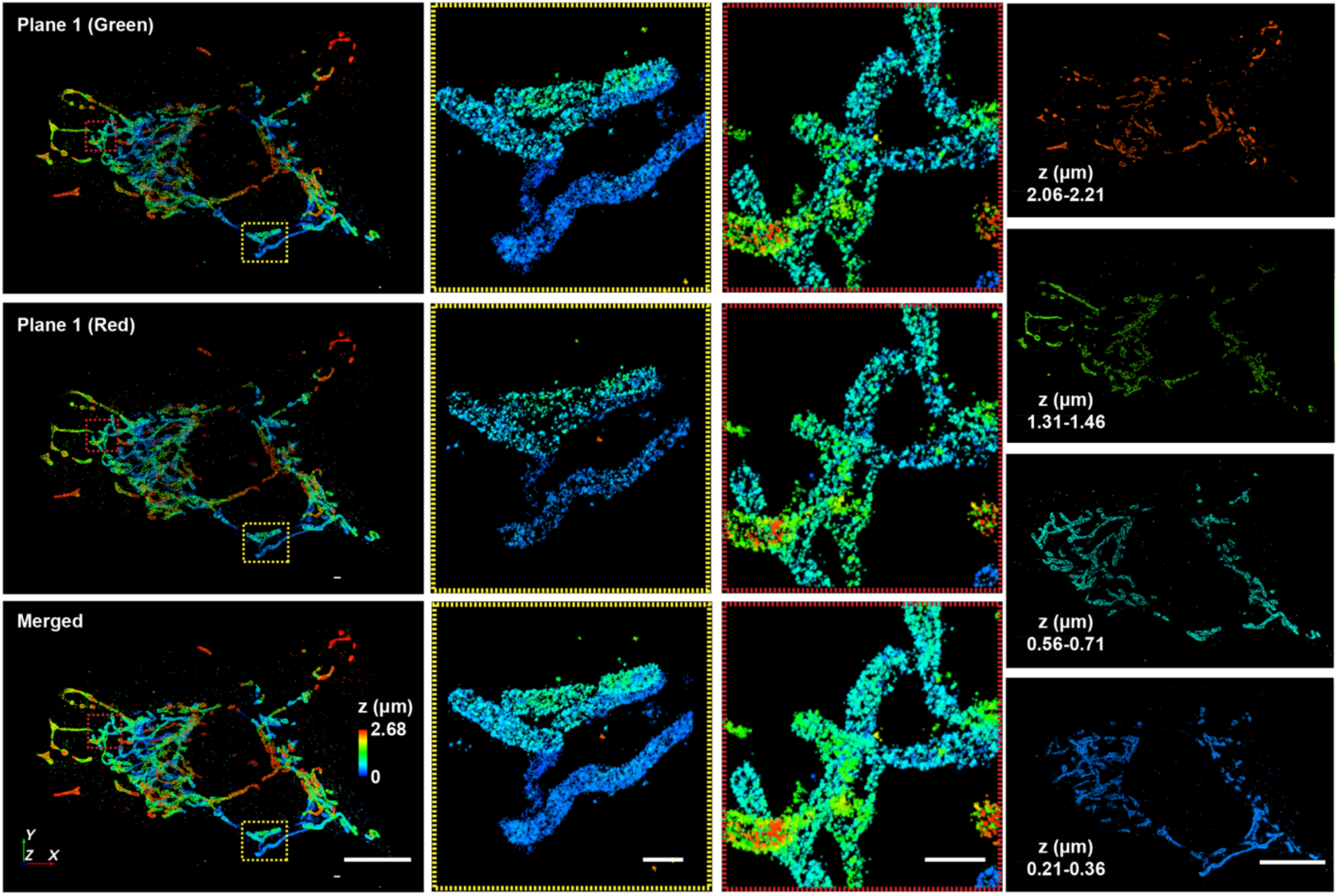
SoLiD-3D improves the precision and speed of 3D single-molecule super-resolution imaging. Reconstructions showing single-target, single-plane imaging of mitochondria (TOMM20) using dual-wavelength excitation and detection for increased speed with retained optical sectioning. The top left panel shows the mitochondria imaged using Cy3B-labeled imager strands in the green channel, the middle left panel shows mitochondria imaged using ATTO 647N-labeled imager strands in the same plane in the red channel, and the bottom left panel shows the merged reconstruction from both channels. The middle columns show insets at the indicated dashed boxes. The right column shows the merged reconstruction at the indicated axial ranges. Scale bars: full field of view: 10 µm, insets: 2 µm, z-slices: 10 µm.

### Parallel whole-cell imaging with improved precision and speed

Whole-cell imaging is typically achieved using wide-field epi-illumination, which degrades the contrast and hence the achievable precision. Simultaneous whole-cell 3D imaging can be achieved using engineered PSFs with long axial range covering the full thickness of the cell, such as DH-PSFs with ∼6 µm axial range (DH-6), but these PSFs suffer from an inherent tradeoff between axial range and achievable precision^31^, further degrading the reconstruction quality. To extend the accessible axial imaging range to enable simultaneous whole-cell imaging while allowing for contrast improvement with thin LSs and improved precision through the use of short axial range PSFs, we implemented a biplane modality in which fluorescence was excited by two axially offset LSs and the emission was recorded simultaneously from two axially offset focal planes. In this configuration, the volumetric coverage is extended without loss of precision or acquisition speed. Here, the red detection channel was axially shifted in relation to the green channel by translating the relay lens in its 4f detection path to be positioned several micrometers above the green plane, with a controlled partial overlap of ∼200–300 nm to ensure continuous axial coverage while minimizing redundancy (Supplementary Fig. 6). Under this configuration, emitters located in the lower axial range were detected in the green channel, while emitters in the upper axial range were detected in the red channel (Fig. 1g). Importantly, each axial plane was illuminated by its corresponding LS, maintaining optical sectioning and minimizing crosstalk between planes. In contrast to conventional biplane implementations that rely on a 50:50 beam splitter and consequently halve the photon budget per channel, our spectrally separated approach preserves photon efficiency while maintaining optical sectioning through LS excitation. To demonstrate this configuration, labeled TOMM20 was excited and detected independently in the two axially offset spectral channels. Localizations from the two color channels were then combined in post-processing to generate a whole-cell 3D reconstruction (Fig. 3). Visualizing distinct z planes of the reconstructed volume reveal continuous structural features of the mitochondria spanning from the lower to the upper detection range, confirming that the two planes seamlessly bridge the full depth of the sample. The merged reconstruction exhibited localization density across an extended axial range of approximately 6 µm, effectively doubling both the coverage and acquisition speed achievable with a single DH-3 PSF channel, and without the precision tradeoff of using a PSF with 6 µm axial range. The spatial resolution was quantified using FRC analysis, yielding resolutions of 40/59/62 nm in the xy/xz/yz planes, respectively (Supplementary Fig. 8).

**Figure 3.**
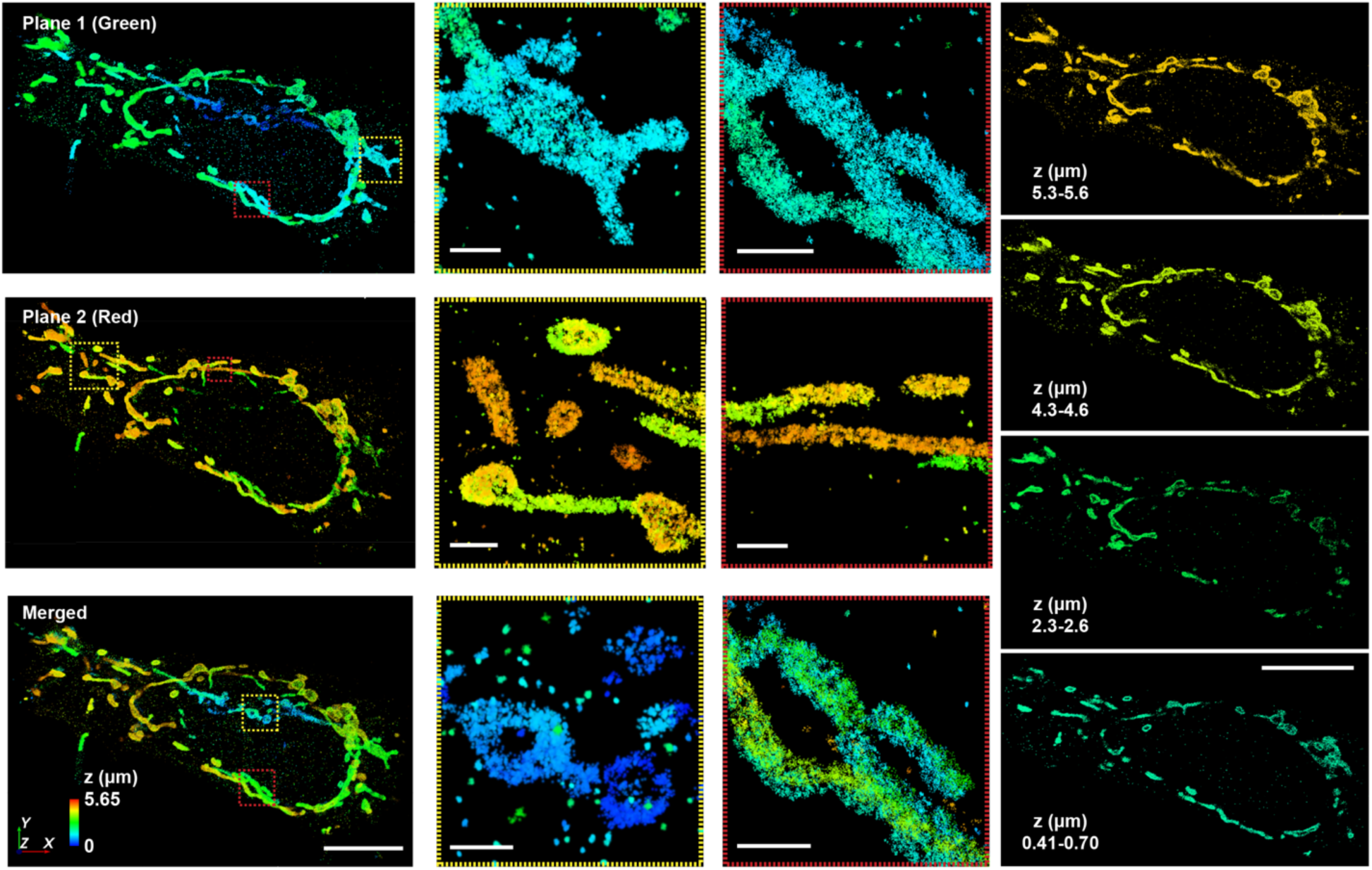
Whole-cell single-molecule super-resolution imaging using PSF-engineering enhanced biplane detection for extended axial coverage. Whole-cell super-resolution reconstructions of mitochondria (TOMM20) was achieved using dual-wavelength, axially offset light sheet (LS) excitation and spectrally separated biplane detection with engineered PSFs in each path. The top left panel shows the mitochondria imaged using Cy3B-labeled imager strands in the green channel at a lower axial plane, the middle left panel shows mitochondria imaged using ATTO 647N-labeled imager strands in an axially offset plane in the red channel, and the bottom left panel shows the merged reconstruction of both planes, providing continuous axial coverage across an extended ∼6 µm range. The middle columns show insets at the indicated dashed boxes. The right column shows the merged reconstruction at the indicated axial ranges. Scale bars: full field of view: 10 µm, insets: 1 µm, z-slices: 10 µm.

### Simultaneous dual-target imaging

The SoLiD-3D platform easily enables multiplexed imaging, which is advantageous for studying multiple targets, such as cellular organelles, at the nanoscale. To demonstrate the capability of the system to perform simultaneous two-target 3D SMLM and to resolve the spatial relationship between subcellular structures throughout cells, we simultaneously imaged mitochondria and the nuclear lamina using DNA-PAINT imaging. TOMM20 was labeled with ATTO 647N imager strands, illuminated with the 647 nm laser, and detected in the red channel, while lamin A/C was labeled with Cy3B imager strands, excited with the 560 nm laser, and detected in the green channel. The emission signals from the two targets were spectrally separated and recorded simultaneously on the same camera. This enabled efficient dual-color acquisition without temporal offset between channels. In the reconstructed volumes, mitochondria are observed wrapping around the nucleus, illustrating the spatial relationship between the mitochondrial network and the nuclear lamina (Fig. 4). The spatial resolution was quantified using FRC analysis, yielding resolutions of 55/60/60 nm in the xy/xz/yz planes for mitochondria, respectively, and 60/61/60 nm in the xy/xz/yz planes for the nuclear lamina, respectively (Supplementary Fig. 9).

**Figure 4.**
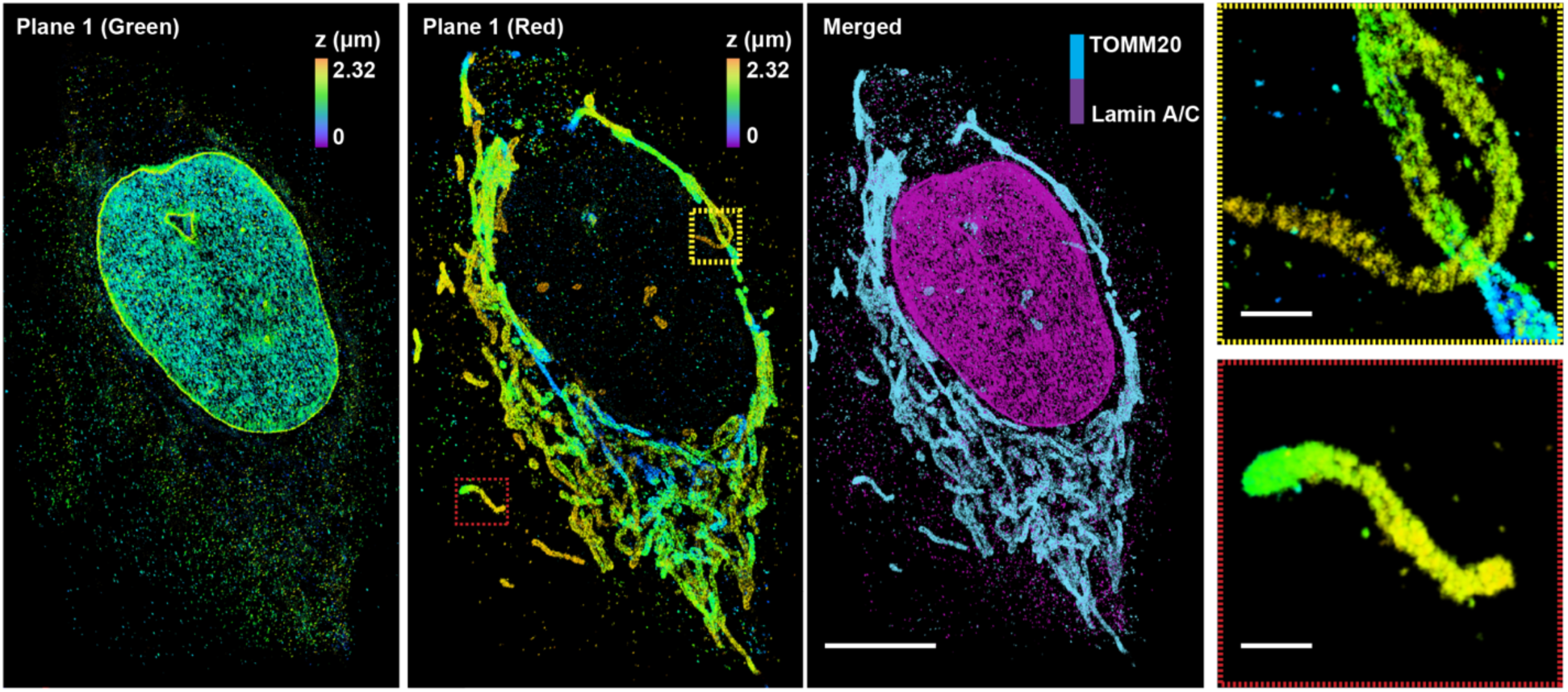
Simultaneous two-target imaging of mitochondria and the nuclear lamina. Reconstructions showing the spatial organization of the nuclear lamina (lamin A/C) and mitochondria (TOMM20). Fluorescence from the two targets was detected using spectrally separated channels, where the green channel corresponds to nuclear lamina and the red channel corresponds to mitochondria. Insets show magnified views of the indicated boxes highlighting the fine structural organization of mitochondria. Scale bars: full field of view: 10 µm, insets: 1 µm.

### The Hummus PSF offers a small spatial footprint with retained axial sensitivity

The DH-PSF encodes axial position through the relative rotation of spatially separated lobes, which enables robust *z*-localization but inherently increases the PSF footprint and limits achievable emitter density^31^. Extending the axial range of the DH-PSF design further intensifies this trade-off, resulting in reduced localization precision and diminished performance under high-background or high-density conditions. To address these limitations, we employed a new type of engineered PSF termed the Hummus-3 PSF, which was optimized using the neural network DeepSTORM3D (DS3D)^36^ (Fig. 5a). Rather than explicitly enforcing geometric constraints on the PSF shape, the design process optimized the shape of the PSF jointly with the decoding neural-net, in an end-to-end fashion, across the axial range of interest, yielding a PSF that concentrates signal into a compact region while retaining high sensitivity to axial position. As a result, the Hummus-3 PSF encodes 3D information with high axial sensitivity without relying on widely separated lateral profiles, thereby enabling further improved performance in high-density imaging regimes.

**Figure 5.**
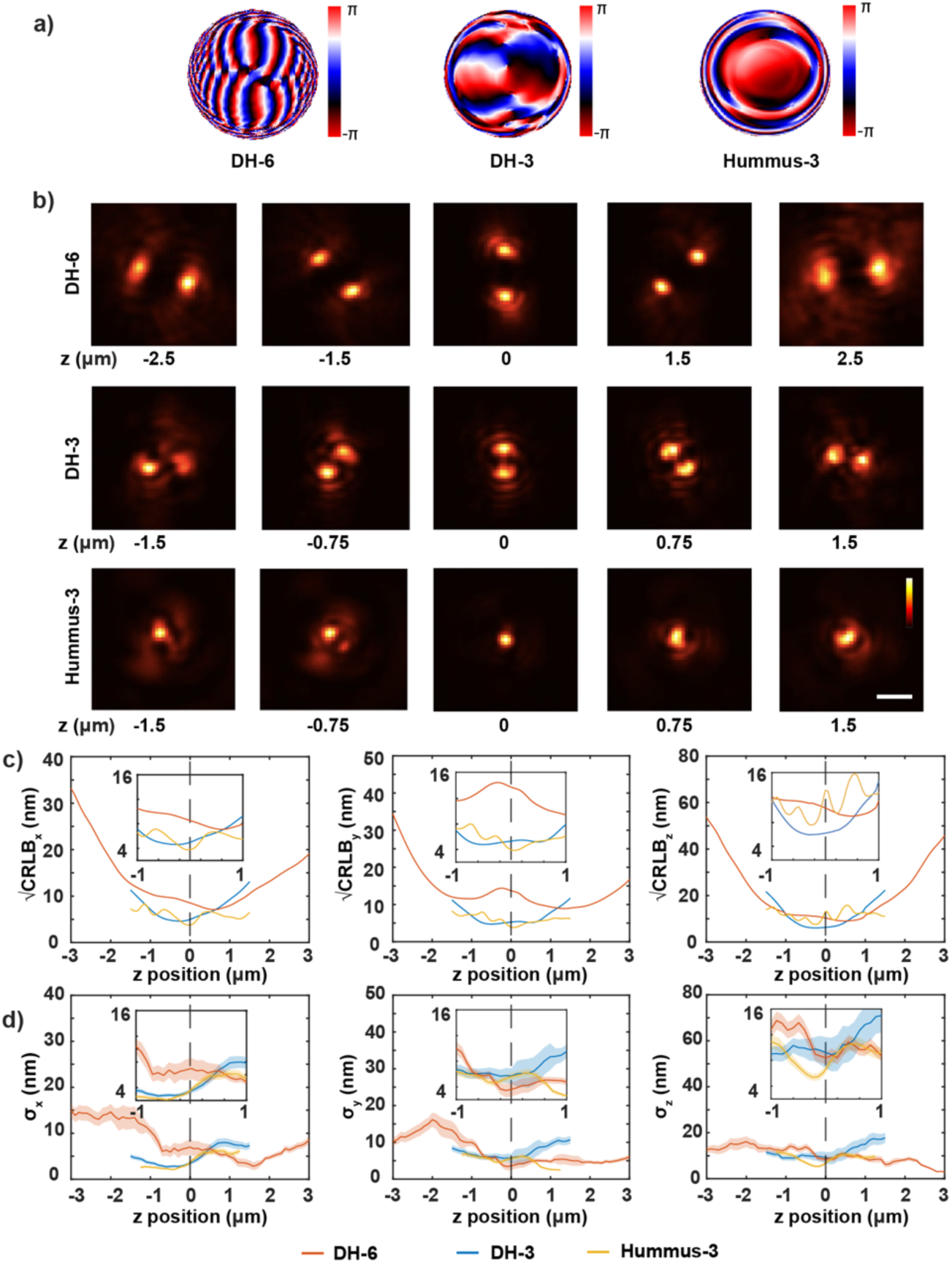
Simulated and experimental comparison of localization precision for engineered PSFs. (a) Three phase masks were used and compared in this study: a double-helix (DH) phase mask designed for a 6 µm axial range (DH-6), a DH phase mask designed for a 3 µm axial range (DH-3), and a newly optimized Hummus phase mask designed for a 3 µm axial range (Hummus-3). The DH and Hummus phase patterns were generated from experimentally obtained z-scans and implemented using the VIPR framework. (b) Simulated PSFs for the DH-3, Hummus-3, and DH-6 phase masks, illustrating the relative PSF footprints and axial encoding characteristics. Scale bar: 2 µm. (c) Numerically calculated localization precision in the x, y, and z dimensions, defined as the square root of the Cramér–Rao lower bound (CRLB¹ᐟ²), computed from the simulated PSFs. Simulations used 6,000 signal photons and a mean background of 10 photons per pixel. (d) Experimental localization precision obtained from repetitive localization of fluorescent beads imaged with the three phase masks. All experimental measurements were acquired under the same conditions. Shaded areas represent the standard deviation of 100 measurements for each z.

The theoretical performance of the Hummus-3 PSF was quantified using the Cramér–Rao lower bound (CRLB), which provides a fundamental limit on localization precision for an unbiased estimator under Poisson noise. CRLB calculations based on simulated PSFs revealed that Hummus-3 PSF achieves localization precision comparable with that of the short-range DH-3 PSF across all three spatial dimensions, while significantly outperforming the long-range DH-6 PSF (Fig. 5b-c). Experimental measurements using sub-diffraction fluorescent beads corroborated these theoretical predictions (Fig. 5d). Localization precision, quantified from repeated localizations of individual beads, showed that the Hummus-3 PSF provides localization precision comparable to that of DH-3, while outperforming the DH-6 across the tested axial range (Fig. 5d). Together, both the simulated and experimental results establish that the Hummus-3 PSF offers high axial sensitivity with a small spatial footprint, making it particularly well suited for high-density, high-speed 3D SMLM.

### The compact footprint of the Hummus PSF enables high-density single-molecule imaging

Having established that the Hummus-3 PSF provides localization precision comparable to or better than conventional DH-PSFs over a 3 µm axial range, we next evaluated how the different PSFs perform at high emitter density. Specifically, we assessed reconstruction performance under increasing emitter densities using simulated datasets with neural network-based localization software AutoDS3D^42^ across all PSF designs (Fig. 6). Detection accuracy, quantified using the Jaccard index, was found to decrease with increasing emitter density for all methods (Fig. 6a-g), consistent with increased PSF overlap and consequent ambiguity in localized emitter position. However, the Hummus-3 PSF consistently exhibits higher detection accuracy, particularly in the high-density regime where overlapping PSFs have greater occurrence. Our data also demonstrates that the Hummus-3 PSF achieves a higher true positive rate and lower miss rate compared to both the DH-3 and DH-6 PSFs, while maintaining a comparable or lower false positive burden. Taken together, these results show that the more compact PSF footprint reduces ambiguity in emitter separation and improves localization robustness.

**Figure 6.**
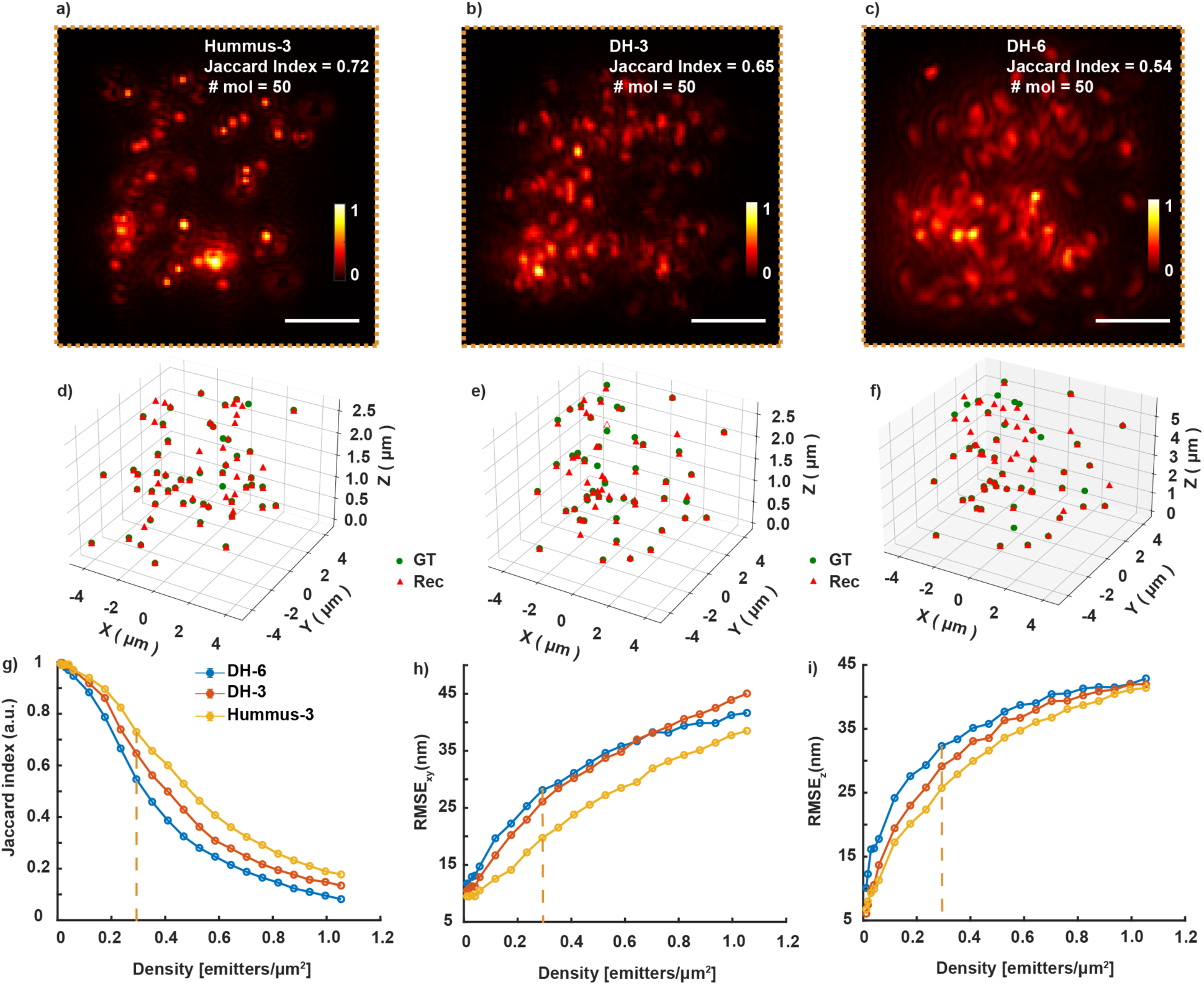
Comparative performance analysis of engineered PSFs under varying emitter densities. (a-c) Representative simulated images of 50 recognized molecules using the DH-6, DH-3, and Hummus-3 PSFs. The resulting Jaccard index is indicated for each PSF at an emitter density of 0.3 emitters/µm². Scale bars: 2 µm. (d-f) 3D scatter plots comparing ground truth (GT, green circles) with reconstructed (Rec, red triangles) molecule positions for the cases shown in (a-c), for DH-6 (J = 0.54, 61 detections, TP = 39, FP = 22, and FN = 11), for DH-3 (J = 0.65, 57 detections, TP = 42, FP = 15, and FN = 8), and for Hummus-3 (J = 0.72, 64 detections, TP = 48, FP = 16, and FN = 3). (g-i) Quantitative metrics as a function of emitter density (emitters/µm²): (g) Jaccard index (detection accuracy), and root mean square error (RMSE) in the (h) lateral plane and (c) axial dimension. The vertical dashed orange line indicates a reference density of 0.3 emitters/µm². Abbreviations: J: Jaccard index, TP: true positives, FP: false positives, FN: false negatives.

Consistent with detection performance, quantification of localization errors also reflects the advantage of the compact PSF design. Both lateral root mean square error (RMSE) and axial RMSE increase with emitter density across all methods; however, the rate of degradation is significantly reduced for the Hummus-3 PSF compared to the DH-3 and DH-6 PSFs (Fig. 6h,i). In particular, axial localization error, which is most sensitive to PSF overlap, remains substantially lower for Hummus-3, highlighting its improved robustness in dense emitter conditions.

We further quantitatively compared the packing density achievable with the different PSFs. The compact spatial footprint of the Hummus-3 PSF enabled substantially higher emitter densities compared with DH-3 and DH-6 (Supplementary Fig. 10a). Within a single 128 × 128 pixels imaging region, the Hummus-3 PSF accommodated up to 272 emitters, compared with 134 emitters for DH-3 PSF and 94 emitters for DH-6 PSF. Thus, relative to the commonly used DH-3 configuration, the Hummus-3 PSF provided more than a 2-fold increase in emitter density per frame while preserving high axial sensitivity (Supplementary Fig. 10a). The lower packing efficiency of the DH-6 PSF resulted from its larger lobe separation, which increased the effective PSF footprint and reduced the number of simultaneously resolvable emitters.

To further quantify spatial efficiency, we measured the foreground pixel occupancy of each PSF across the calibrated axial range. The Hummus-3 PSF consistently occupied approximately 2-fold fewer pixels than the DH-3 PSF throughout the imaging volume (Supplementary Fig. 10b), confirming its compact footprint. This reduced pixel occupancy directly translates to lower overlap-induced fitting errors and increased localization throughput under high-density imaging conditions.

### The Hummus PSF provides high-fidelity 3D super-resolution imaging in cellular environments

Finally, we demonstrated the practical utility of the Hummus-3 PSF for 3D super-resolution imaging in cells at high emitter densities by imaging of mitochondria. These experiments provide a proof-of-principle application of the Hummus-3 PSF in a biologically relevant sample, highlighting its capability for volumetric single-molecule imaging in complex intracellular environments (Supplementary Video 3). The higher emitter densities enabled by the Hummus-3 PSF allowed dense sampling of the mitochondria, resulting in robust reconstruction under simultaneous dual-wavelength excitation and two-color detection of the same target with improved temporal resolution (Fig. 7a-d). The green (Fig. 7b) and red (Fig. 7c) channels independently resolved mitochondrial morphology with high spatial consistency, while the merged reconstruction (Fig. 7d) provided a more continuous and complete representation of the mitochondrial network compared to either channel alone, highlighting improved structural sampling enabled by increased localization density. Further analysis of zoomed-in regions of interest (Fig. 7d) revealed that the merged reconstruction enhances continuity of mitochondrial filaments, reducing fragmentation observed in the individual channels and improving visualization of fine structural details. The spatial resolution was quantified using FRC analysis, yielding resolutions of 73/80/81 nm in the xy/xz/yz planes, further demonstrating the robustness of the Hummus-3 PSF for high-fidelity 3D super-resolution imaging in densely labeled cell environments (Supplementary Fig. 11).

**Figure 7.**
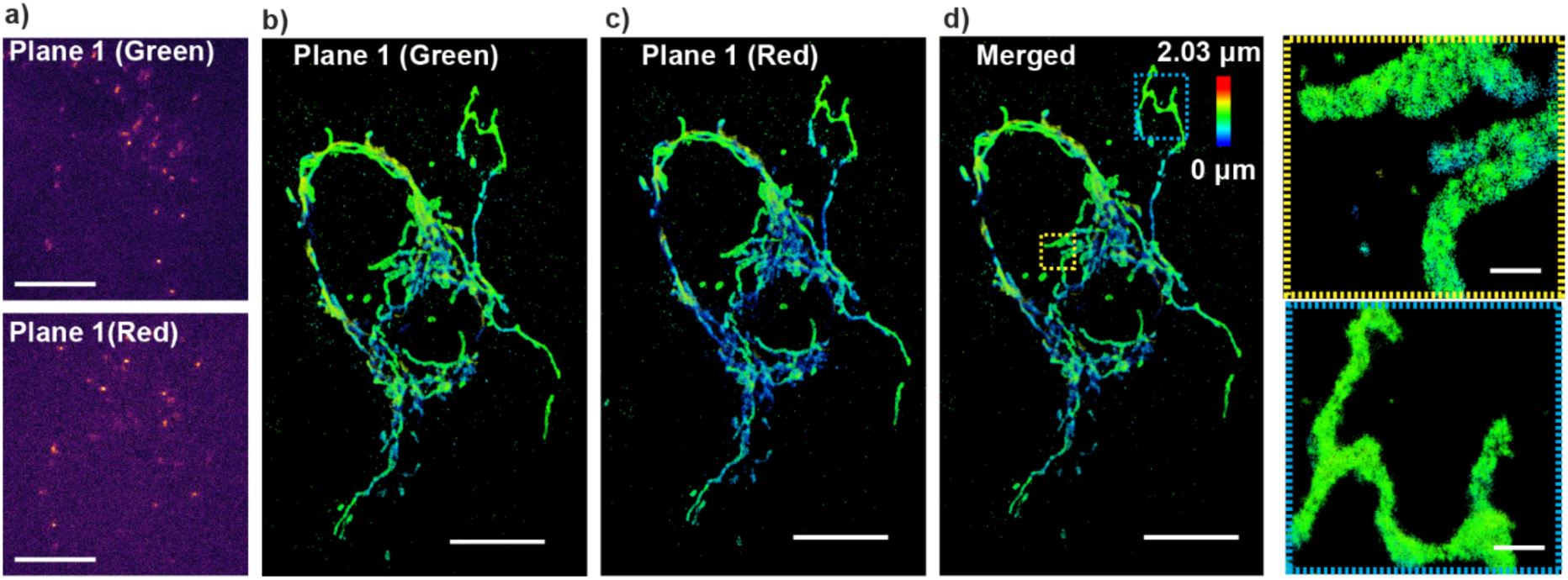
High-density 3D single-molecule super-resolution imaging enabled by the Hummus-3 PSF with compact footprint. (a) Representative single-molecule images acquired using the Hummus-3 PSF. Scale bars: 5 µm. (b-d) 3D single-molecule super-resolution reconstructions of mitochondria (TOMM20) acquired using the Hummus-3 PSF under dual-wavelength excitation, shown separately for the (b) green channel, (c) red channel, and (d) merged. The insets show zoomed-in regions at the indicated dashed boxes of the merged reconstruction. Scale bars: full field of view: 10 µm, insets: 1 µm.

## Discussion

Three-dimensional SMLM faces trade-offs between photon efficiency, background suppression, axial range, and achievable emitter density. In this work, we introduced SoLiD-3D, a LS-based, spectrally separated biplane imaging platform that overcomes these limitations by integrating dual-wavelength excitation, photon-efficient detection, and dual-channel PSF engineering within a compact and flexible optical architecture. A central advantage of SoLiD-3D is the replacement of intensity-based beam splitting with spectral separation, enabling simultaneous biplane detection without compromising the photon budget. When combined with dual-wavelength LS excitation, this approach further suppresses out-of-focus background and minimizes crosstalk between detection planes. The resulting improvement in the SBR directly translates into enhanced localization precision. The measured reductions in fluorescence background in cell samples using LS excitation confirm the effectiveness of optical sectioning in dense cellular environments. Importantly, these improvements were achieved without compromising signal photon counts, illustrating the compatibility of LS illumination with high-NA single-objective detection.

Beyond improving localization precision, SoLiD-3D enables switchable imaging modalities and parallelization of localization throughput by exploiting spectral degrees of freedom. Simultaneous dual-color imaging of the same biological target effectively doubles localization density per frame without necessitating increased emitter overlap or reducing either channel’s photon counts. This strategy contrasts with traditional multiplexing approaches that rely on temporal separation or emission splitting, both of which impose barriers towards improved imaging speed or localization precision. The nanoscale resolution confirmed by FRC analysis demonstrates that spectral parallelization can be used not only to accelerate acquisition, but also to maintain high structural fidelity across extended volumes. This capability is particularly valuable for whole-cell volumetric imaging, where acquisition times are often constrained by fluorophore photophysics, labeling density, or analysis software. By decoupling localization density from photon loss, SoLiD-3D provides a scalable route toward faster, high-quality 3D reconstructions. SoLiD-3D further extends this volumetric imaging capability through spectrally encoded biplane detection, in which two focal planes with tunable axial offset are imaged simultaneously in separate detection channels. Precise calibration of relay-lens translation to axial displacement ensures controlled plane separation and predictable overlap, enabling continuous axial coverage over ∼6 µm without gaps. Compared to single-plane imaging with extended-range PSFs^34^, this approach preserves localization precision while mitigating the loss of sensitivity that typically accompanies larger axial ranges. The complementary sampling observed in the overlap region between planes highlights the robustness of this configuration and demonstrates its suitability for whole-cell imaging. Importantly, because the biplane modality is implemented spectrally rather than by intensity splitting, it retains the photon efficiency advantages of the single-plane dual-color configuration. Together, these features render SoLiD-3D a flexible platform capable of improved axial range, speed, and localization density without compromising signal quality.

While optical strategies can improve background suppression and photon efficiency, localization throughput in SMLM is ultimately constrained by the PSF footprint and emitter overlap, such that even advanced neural network-based localization methods remain fundamentally limited at high emitter densities. To address this limitation, we introduced the Hummus-3 PSF, a compact PSF design derived by end-to-end optimization rather than explicit geometric constraints. By redistributing Fisher information more efficiently across the axial range, the Hummus-3 PSF achieves high axial sensitivity while maintaining a substantially reduced spatial footprint compared to conventional DH-PSFs. Both CRLB analysis and experimental measurements confirmed that the Hummus-3 PSF provides localization precision comparable to or better than short-range DH-PSFs, while significantly outperforming long-range DH-PSF designs that suffer from expanded footprints and degraded axial precision. Importantly, the compactness of the Hummus-3 PSF enables more than a 2-fold increase in achievable emitter density per frame relative to the DH-3 PSF, directly translating into higher localization throughput under high-density conditions. This capability is particularly advantageous for fast volumetric imaging of complex cellular structures, where high labeling densities and limited acquisition times demand efficient use of detected photons.

By integrating dual-LS excitation, spectrally separated and parallelized detection, PSF-engineering enhanced biplane volumetric imaging, and compact PSF engineering, SoLiD-3D addresses multiple, traditionally competing optimizations in 3D SMLM within a single, unified optical framework. The modular nature of the platform allows individual components, such as spectral parallelization or compact PSFs, to be deployed independently or in combination, depending on experimental priorities. While the current implementation focuses on dual-color imaging, the underlying principles readily extend to additional spectral channels or alternative PSF designs optimized for specific axial ranges or imaging speeds. More broadly, this work illustrates how combining optical architecture with PSF design can shift fundamental limits in localization microscopy and increase temporal resolution. By maximizing photon efficiency and minimizing spatial redundancy, SoLiD-3D enables rapid, high-density, whole-cell 3D imaging with nanoscale precision, establishing a powerful framework for next-generation volumetric microscopy. This capability paves the way for real-time visualization of living cells, continuous tracking of nanoscale structural remodeling, and comprehensive mapping of dynamic molecular interactions across entire cellular volumes. In doing so, SoLiD-3D opens new opportunities to interrogate complex, heterogeneous biological systems, bridging the gap between static super-resolution and true spatiotemporal nanoscale biology.

## Methods

### SoLiD-3D optical platform

The SoLiD-3D microscope integrates dual-color, single-objective LS illumination with tunable biplane detection with engineered PSFs in the emission pathway (Fig. 1, Supplementary Fig. 1). The system is configured to allow rapid switching between LS and conventional epi-illumination modes. In addition, the detection scheme is modular, enabling straightforward switching between biplane detection and single-plane imaging within the same optical platform.

Two continuous-wave visible fiber lasers at 560 nm and 647 nm (1 W each, 2RU-VFL, MPB Communications) were used for excitation. The laser beams were spectrally filtered (FF01-554/23-25, Semrock; FF01-643/20-25, Semrock) and circularly polarized (560 nm: LPVISC050-MP2 polarizer, Thorlabs, and Z-10-A-.250-B-556 quarter-wave plate, Tower Optical; 647 nm: LPVISC050-MP2 polarizer, Thorlabs, and Z-10-A-.250-B-647 quarter-wave plate, Tower Optical). Switching between lasers was controlled with shutters (VS14S2Z1, Vincent Associates Uniblitz) connected to a shutter control box (VDM-D3, Vincent Associates Uniblitz). Each beam was then expanded 6-fold and collimated using telescopes (AC254-025-A-ML, focal length of 25.4 mm and AC254-150-A-ML, focal length of 150 mm, Thorlabs). The expanded beams were then coaligned using a dichroic mirror (T590lpxr, 26 mm × 38 mm, Chroma Technology). The combined beam was elevated to the height of the microscope illumination input port (Ti2-E, Nikon inverted microscope) using a periscope assembly and subsequently directed toward a flip mirror (TRF90, Thorlabs), which enabled rapid switching between epi-illumination and LS illumination modes.

In the epi-illumination path, the beam was further expanded using a second beam expander (AC254-075-A-ML and AC254-300-A-ML, Thorlabs) and routed via an additional flip mirror (TRF290, Thorlabs) to a Köhler lens (ACT508-400-A-ML, Thorlabs). This lens focused the beam onto the back focal plane of a high-NA TIRF objective (CFI SR HP Apo TIRF 100x, NA 1.49 OIL, WD 0.12 mm, FOV 22 mm, Nikon), enabling widefield epi-illumination.

For LS illumination, the spectrally coaligned beam was split again into its constituent wavelengths using a dichroic mirror (T590lpxr, 26 mm × 38 mm, Chroma). The 560 nm and 647 nm beams were subsequently processed along independent but symmetric optical paths until recombination. Each beam was first demagnified using a telescope composed of achromatic lenses (AC254-150-A-ML and AC254-075-A-ML, Thorlabs). The 560 nm beam was then passed through a cylindrical lens (ACY254-150-A, Thorlabs) to generate a one-dimensional focus, forming a LS that was directed onto an x-axis galvanometric mirror (G1, GVS211, Thorlabs). The LS thickness was further reduced using an additional demagnifying telescope (AC254-150-A-ML and AC254-075-A-ML, Thorlabs) optimizing the LS thickness to match the axial range of the engineered PSFs and to minimize out-of-focus excitation. Following this, the beam was relayed onto a y-axis galvanometric mirror (G2, GVS211, Thorlabs), which was optically conjugated to G1. The beam was then further demagnified using the same lens combination and imaged onto a focus-tunable lens (EL-30-10, focal length -80 mm to +1000 mm, Optotune), providing rapid positioning of the LS focus. The tunable lens operated under a built-in closed-loop feedback system to maintain a stable focal length during operation. An identical optical configuration was implemented for the 647 nm beam, using a separate pair of galvanometric mirrors (G3 and G4) and identical relay optics. The two spectrally separated LS excitation pathways were maintained fully independent of one another up to after the tunable lenses. To ensure matched optical path lengths, a unit magnifier (both AC254-050-C-ML, Thorlabs) was introduced into the 647 nm excitation path. After the tunable lenses, the 560 nm and 647 nm beams were recombined using a dichroic mirror (T590lpxr, 26 mm × 38 mm, Chroma). The combined beam was subsequently relayed through a 4f system comprising 300 mm (AC254-300-A-ML, Thorlabs) and 400 mm (ACT508-400-A-ML, Thorlabs) focal length lenses. A dithering mirror (SP30Y-AG, Thorlabs; GPWR15 Galvo Power Supply, Thorlabs; CBLS3F Galvo System Cable Set, Thorlabs) was positioned at the Fourier plane of a 4f relay system consisting of a lens (AC254-300-A-ML, Thorlabs) placed 300 mm from the tunable lens and a Köhler lens (ACT508-400-A-ML, Thorlabs) positioned 400 mm from the back aperture of the objective lens. Rapid oscillation of the dithering mirror with a function generator (15 MHz DDS Signal Generator/Counter, Koolertron) in the plane of the LSs effectively averaged out shadowing and striping artifacts arising from reflective structures or inhomogeneities within the sample. For all experiments, the oscillation frequency was fixed at 100 Hz with a driving amplitude of 0.1 V. All galvanometric mirrors (G1–G4) were precisely aligned and optically conjugated to the back focal plane of the objective, enabling accurate translation of the LSs in the lateral (*x* and *y*) directions. G1 and G3 were used to laterally align the 560 nm and 647 nm LSs along the x-direction, while G2 and G4 controlled alignment along the axial direction, enabling either coincident illumination or axial separation for biplane excitation.

Fluorescence emitted from the sample was collected by the same objective lens (CFI SR HP Apo TIRF 100x, NA 1.49 OIL, WD 0.12 mm, FOV 22 mm, Nikon) used for excitation and imaged onto the microscope image plane by the internal tube lens. An adjustable iris placed at this intermediate image plane (103 mm distance from the microscope body) was used to control the size of the field of view that was relayed onto the camera sensor, allowing two axially co-aligned or separated image planes to be captured simultaneously on a single sCMOS camera (Prime 95B, Teledyne Vision Solutions). For detection, the emission was separated into two spectral channels using a dichroic mirror (T590lpxr, 26 mm × 38 mm, Chroma Technology) placed after the first lens (AC254-080-A-ML, Thorlabs) of a 4f relay system. In the green detection path, the second 4f lens (AC254-080-A-ML, Thorlabs) was kept fixed, while in the red detection path the corresponding lens was mounted on a three-axis translation stage (PT1, Thorlabs, USA). This configuration enabled controlled axial displacement between the two image planes, forming a biplane detection system. Engineered phase masks (green: DH1-580, or DH6R-580, red: DH1-670, DH6R-670 respectively, all Double Helix Optics, Inc. or Hummus-3) were placed at the Fourier plane of each detection channel to encode axial position information into the PSF. This detection architecture supports multiple imaging modes, including single-target imaging with co-aligned LSs, dual-color imaging of two molecular targets, and extended-depth imaging using axially separated biplane detection and spectrally separated illumination and detection.

### Volumetric imaging characterization and calibration

To quantify the volumetric performance of SoLiD-3D, we first characterized the performance of the DH-3 phase mask. Fluorescent beads (T7280, TetraSpeck, 0.2 µm, Invitrogen) were immobilized on a coverslip and imaged over an axial range of 3 µm using a piezo-controlled nano-positioning stage (Nano-LPS series, Mad City Labs GmbH, Kloten, Switzerland). Images were acquired at 100 nm axial increments using the sCMOS camera controlled via Micro-Manager^43^, with custom acquisition routines implemented in MATLAB. At each z position, the angular orientation of the DH-PSF lobes was quantified using EasyDHPSF^44^. Consistent with the design of the DH-3 phase mask, the two PSF lobes exhibited a robust and continuous rotation as a function of axial displacement (Supplementary Fig. 6a). This angular rotation provided a one-to-one mapping between lobe orientation and emitter z position across the usable axial range. A calibration curve was constructed by fitting the measured lobe angles to the known stage positions, establishing a precise angular-to-axial conversion using EasyDHPSF^39^. This calibration serves as the reference framework for all subsequent axial localizations and for alignment of the two detection planes in the biplane configuration. Using this angular calibration, we next established the relationship between mechanical translation in the detection relay optics and the corresponding axial displacement at the sample plane. The focal plane was determined so that the DH-3 PSF was first positioned at a defined angular orientation corresponding to a known axial coordinate, typically near the upper end of the DH-3 encoding range (∼1.4 µm), and an image was acquired (Fig. 1g). The relay lens was then translated in discrete steps of 25 µm, and at each position ten images were recorded to ensure statistical robustness. Each acquired PSF image was analyzed using EasyDHPSF, and the extracted lobe angles were converted to axial positions using the previously established DH-3 calibration curve (Supplementary Fig. 6b). This procedure generated a precise mapping between relay lens displacement and effective axial shift at the sample plane (Supplementary Fig. 6b). From this calibration, we determined that a 1 mm translation of the relay lens corresponds to an approximately 100 nm axial displacement in the object plane. This conversion factor enables deterministic and reproducible positioning of the second detection plane for biplane imaging.

Having established the axial and relay calibration, we evaluated the full biplane configuration by simultaneously imaging single fluorescent beads (T7280, TetraSpeck, 0.2 µm, Invitrogen) under dual excitation at 560 nm and 647 nm. To generate two axially separated detection planes, the second relay lens in the red detection path was translated by approximately 2.5 µm at the sample plane, as determined from the lens-to-z calibration. This offset was selected to produce two partially overlapping axial detection volumes with an overlap of approximately 200–300 nm, ensuring continuous coverage while minimizing redundancy (Fig. 1h). The axial performance of both channels was then assessed across an extended imaging range of ∼5–6 µm. Within the 0–3 µm range, the DH-3 in the green detection channel remained well-defined and fully resolvable, while the red channel signal was negligible due to the intentional axial offset (Fig. 1g). As the emitter was displaced beyond ∼2.7 µm, the DH-3 PSF in the green channel approached the limit of its encoding range and gradually lost localizability. Concurrently, in the red channel, a clearly detectable DH-3 PSF could be localized, indicating that the emitter had entered the axial range of the red detection path (Fig. 1g).

### Microfabrication process for reflective microfluidic inserts

The micro-optics incorporated microfluidic chips were fabricated as described previously^10,20,45^. In brief, the inserts were designed in AutoCAD (Autodesk) and prepared for 3D nanoprinting using Describe (Nanoscribe GmbH & Co.KG) with the 10× Silicon Shell recipe (0.3 µm slicing and hatching distances, -6° hatching angle), with block parameters optimized to eliminate stitching artifacts. Inserts with widths of 300 µm and lengths of 2.5–4 mm were fabricated with heights of 105 µm for mammalian cell imaging and LS characterization and sidewall angles of 39° or 45° with mirror heights of 15–70 µm to generate reflected LS geometries with defined tilt. Structures were printed by two-photon polymerization on fused silica using a Nanoscribe Photonic Professional GT2 (Nanoscribe GmbH & Co.KG) with the IP-Visio resin (Nanoscribe GmbH & Co.KG), followed by development with SU-8 Developer (mr-Dev 600, Kayaku Advanced Materials, Inc.), isopropyl alcohol (MPX18304, Fisher Scientific) rinsing, nitrogen drying, and UV curing (140010, Vacuum UV-Exposure Box, Gie-Tec GmbH). Inserts were selectively metalized by e-beam deposition with silica (200 nm) (EVMSIO21-5D, Kurt J. Lesker), aluminum (350 nm) (EVMAL50EXEB, Kurt J. Lesker), and silica (5 nm) (EVMSIO21-5D, Kurt J. Lesker) to form thermally protected reflective mirrors. PDMS microfluidic bases were fabricated by casting SYLGARD 184 (10:1 elastomer to curing agent) over SU8-100 molds patterned on silicon wafers with channel heights of 100 µm using standard photolithography, followed by curing overnight at room temperature and for 2 h at 65 °C. Metalized inserts were placed into the PDMS channels in the correct orientation for LS reflection, and the PDMS was plasma-bonded (PDC-32G, Harrick Plasma Inc) to cleaned glass coverslips (#1.5, 22 mm × 22 mm, Electron Microscopy Sciences) to form sealed microfluidic devices.

### Hummus phase mask design and fabrication

Phase mask design was carried out using the end-to-end optimization framework implemented in DeepSTORM3D^34^, with the objective of generating an optimized PSF over a 3 µm axial range. Phase masks for the two emission channels were independently optimized with uniform background and a mask pixel size of 20 µm. For fabrication, the optimized continuous phase profiles were converted into height maps for photolithography. Material dispersion of fused silica was used to account for wavelength-dependent phase modulation between the 590 nm and 680 nm emission channels. These height maps were then discretized into 16 final height levels spanning 0–1.5 µm, with 0.1 µm spacing between adjacent levels. The final multi-level phase masks were fabricated by HOLO/OR using a pixel size of 10 µm and four binary fabrication steps to implement the 16-level profiles.

### SoLiD-3D laser beam steering calibration and excitation beam characterization

Calibration of the LS steering was performed by directly imaging the excitation beam in a fluorescent solution containing 1 µM Cy3B- or ATTO 647N-labeled DNA imager strands (Massive Photonics, order number AN2601015). All measurements were carried out in the absence of the reflective insert. The sample was mounted in a standard imaging chamber (8 well, ibidi). Each of the four galvanometric mirrors (G1–G4) was calibrated independently under identical optical conditions (Supplementary Fig. 12). The mirrors were driven using a PCIe-based data acquisition system (BNC-2090A, National Instruments) with analog voltage outputs controlled through custom-written MATLAB codes. For each mirror, the applied control voltage was varied over a maximum range of -0.5 V to +0.5 V in increments of 0.01 V, while keeping all other mirrors at a fixed reference voltage (typically 0 V). At each voltage step, an image of the LS was acquired with an exposure time of 50–100 ms. For each voltage, 5-10 frames were acquired and averaged to improve the signal-to-noise ratio (SNR). The laser power at the output of the fiber port was set to 150 mW and attenuated using a neutral density filter (ND 2.6). For each acquired image, a one-dimensional fluorescence intensity profile was extracted along the direction orthogonal to beam propagation by selecting the row or column corresponding to the maximum signal intensity. The extracted peak positions were plotted as a function of the applied voltage to generate calibration curves for each galvanometric mirror. Linear regression was used to obtain conversion factors (∼ 1.1 µm per 0.01 V), enabling precise mapping between control voltage and lateral or axial displacement of the LS within the sample.

LS characterization was performed under similar conditions by imaging the LS after reflection off of a 45° microfabricated mirror integrated within the microfluidic chip (channel dimensions: 700 µm wide and 1 cm long) filled with fluorescent solution containing 0.5–1 µM Cy3B- or ATTO 647N-labeled DNA imager strands. Using this scheme, the cylindrical lens was rotated in order to collect images of both the thickness and width of the LS for a complete characterization (Supplementary Figs. 2-3). For each configuration, 10-50 frames were acquired and averaged to improve the SNR. Beam profiles were analyzed in ImageJ^46^ by extracting intensity line scans perpendicular to the direction of beam propagation at different positions along the optical axis. Each profile was fit to a Gaussian function to determine the 1/e^2^ beam waist radius of each distribution. The variation of beam thickness as a function of propagation distance was used to estimate parameters such as the beam waist radius, confocal parameter, and axial tunability (Supplementary Figs. 2-5). These measurements provided a quantitative characterization of the LS geometry and verified consistent beam quality across both excitation wavelengths.

### Cell culture

Human osteosarcoma cells (U-2 OS – HTB-96, ATCC) were thawed from -80°C (passage 6), centrifuged, and the supernatant was aspired. The cell pellet was resuspended in 1 ml cell culture media (Dulbecco’s modified Eagle’s medium (DMEM) with 10% fetal bovine serum (FBS) and 1 mM sodium pyruvate, all Gibco) and seeded in a T25 flask (2 ml of culture media and 1 ml of resuspended cells). The cells were incubated for 24 h in 37°C at 5% CO_2_ and 90% humidity (Thermo Scientific Heracell 150i CO2 Incubator, 51-032-871, Fisher Scientific). After 24 h, the cells were washed with PBS to remove cell debris and then kept in cell culture media for 48-72 h in the incubator at 37°C with 5% CO_2_ and 90% humidity. The cells were split and cultured for 2–3 passages before being seeded in microfluidic channels for experiments.

### Cell seeding and fixation

Before cell seeding inside microfluidic channels, the channels were sterilized by rinsing with 70% ethanol (BP82031GAL, Fisher Scientific), followed by nanopure water, and subsequently functionalized with 0.001% fibronectin (F0895, Sigma–Aldrich) prepared in phosphate-buffered saline (PBS) (SH3025601, Fisher Scientific). After seeding cells and allowing them to attach in the microfluidic chip for ∼8 h in the incubator at 37°C with 5% CO_2_ and 90% humidity, cells were washed twice with PBS, fixed with 4% paraformaldehyde (PFA, Electron Microscopy Sciences) in PBS (SH3025601, Fisher Scientific) for 20 min, washed again with PBS, quenched with 10 mM NH₄Cl (Sigma–Aldrich) in PBS for 10 min, and washed once more with PBS. Samples were either stored in PBS at 4°C or immediately immunolabeled.

### Cell labeling for diffraction-limited imaging

Following fixation, cells were washed once with PBS and incubated in 10 mM ammonium chloride (NH₄Cl) in PBS for 10 min to quench residual aldehyde groups. Cells were then permeabilized by three washes with 0.2% (v/v) Triton X-100 in PBS (pH 7.4), with a 5-min incubation during each wash. After permeabilization, samples were blocked in 3% (w/v) bovine serum albumin (BSA Sigma–Aldrich) in PBS for 1 h at room temperature. For immunolabeling, cells were incubated for 2 h at room temperature with rabbit anti-lamin B1 primary antibodies (1:1,000; ab16048, Abcam) diluted in 1% (w/v) BSA in PBS. Excess primary antibodies were removed by three washes with 0.1% (v/v) Triton X-100 in PBS, with a 3-min incubation between washes. Samples were then incubated for 1 h at room temperature, protected from light, with donkey anti-rabbit secondary antibodies conjugated either with CF568 (1:100; 20098-1, Biotium) or Alexa Fluor 647 (1:100, ab150067, Abcam) diluted in 1% (w/v) BSA in PBS. Following secondary labeling, cells were washed five times with 0.1% Triton X-100 in PBS, followed by ten washes with PBS. Finally, samples were kept immersed in PBS to prevent drying prior to diffraction-limited or super-resolution imaging.

### DNA-PAINT sample preparation

For labeling, cells were first permeabilized with 0.1% (v/v) saponin in PBS for 10 min and blocked for 1 h at room temperature in blocking buffer containing 0.05 mg/mL sheared salmon sperm DNA (15632011, Invitrogen, Thermo Fisher Scientific) and 10% (v/v) donkey serum (ab7475, Abcam) in PBS.

For epi- versus LS single-molecule imaging comparison, samples were incubated for 2 h at room temperature with rabbit anti-lamin B1 primary antibodies (1:1,000; ab16048, Abcam) diluted in 1% (w/v) BSA in PBS (SH3025601, Fisher Scientific). Following primary antibody labeling, samples were washed three times with 0.1% Triton-X 100 (Sigma–Aldrich), with a 3-minute incubation period between each wash. Samples were subsequently rinsed with washing buffer (Massive Photonics) prepared by diluting the stock solution from 10× to 1× in nanopure water before use. Anti-rabbit IgG conjugated to docking strand 2 (Massive-AB 2-Plex, Massive photonics, order no AB2601023) were diluted at a concentration of 1:100 in antibody incubation buffer (Massive Photonics) and were allowed to incubate for 1 h at room temperature. After secondary antibody labeling, samples were washed three times with 1× washing buffer (Massive Photonics). Following a final wash with 1× imaging buffer (Massive Photonics), imager strands diluted in imaging buffer were added for imaging. A 1 μM imager strand stock solution was diluted to 1 nM.

For multi-target DNA-PAINT imaging, samples were incubated with rabbit anti-TOMM20 primary antibodies (ab186735, Abcam) diluted at a concentration of 1:200 and mouse anti-lamin A/C primary antibodies (sc-376248, Santa Cruz Biotechnology) diluted at a concentration of 1:100 in blocking buffer for 1 h at room temperature or overnight at 4°C. Following labeling with primary antibodies, samples were washed three times for 5 min with PBS and once with 1× washing buffer (Massive Photonics). The sample was then labeled using the Massive Photonics DNA-PAINT secondary antibody kit (Massive-AB 2-Plex, Massive photonics, order no AB2601023). Anti-rabbit IgG conjugated to docking strand 2 and anti-mouse IgG conjugated to docking strand 1 were each diluted at a concentration of 1:100 in antibody incubation buffer (Massive Photonics) and were allowed to incubate for 1 h at room temperature. Samples were then washed once with PBS for 7 min, followed by three 7 min washes with 1× washing buffer, and a final 5 min wash with PBS. Labeled samples were stored in PBS at 4°C until imaging. For DNA-PAINT imaging, samples were incubated in imaging buffer containing 1 nM imager strands prepared from 1 µM stock solutions. In multi-target experiments, imager 1 conjugated to Cy3B was used to image docking strand 1 (corresponding to the mouse secondary antibodies used for lamin A/C imaging), while imager 2 conjugated to ATTO 647N was used to image docking strand 2 (corresponding to the rabbit secondary antibodies used for TOMM20 imaging) enabling spectrally separated two-color DNA-PAINT imaging.

For two-color single-target same-plane and biplane imaging, TOMM20 was immunolabeled as above and imaged using Cy3B-labeled and ATTO 647N-labeled imager 2 strands. Identical imager–docking strand pairs (imager 2) were used in both spectral channels to ensure consistent binding kinetics and localization performance across detection paths.

### Fiducial bead preparation

Calibration images of the PSFs were generated using fluorescent fiducial beads (T7280, TetraSpeck, 0.2 μm diameter; Invitrogen) for both 560 nm and 647 nm excitation. The fiducial beads were spin-coated onto coverslips in a 1% (w/v) poly(vinyl alcohol) (PVA; Mowiol 488, #17951, Polysciences Inc.) solution prepared in nanopure water.

For drift correction of single-molecule data, fluorescent beads (T7280, TetraSpeck, 0.2 µm, Invitrogen) were introduced into the microfluidic channel prior to imaging at a low concentration (dilution of 10⁶-10⁷) to minimize interference with single-molecule detection.

### Single-molecule super-resolution imaging data acquisition

Fluorescence was simultaneously recorded in two spatially separated subregions of the sCMOS camera (full camera frame size was 2048 × 2048 pixels), each with an effective size of approximately 512 × 512 pixels. With a calibrated pixel size of 108 nm, each subregion corresponded to a field of view of approximately 55 × 55 μm². The camera conversion gain was 0.56.

For comparison of single-molecule imaging performance under epi- and LS illumination, 5,000 frames were acquired with an exposure time of 100 ms. Image acquisition was performed using Micro-Manager (micro-manager-2.0) to control the sCMOS camera. To enable 3D single-molecule localization and to quantitatively compare background levels between epi- and LS illumination, a DH phase mask (DH-3, Double Helix Optics, Inc.) was placed in the Fourier plane of the detection paths. The acquired data were analyzed using the easy-DHPSF software^44^.

For single-target single-plane DNA-PAINT imaging of mitochondria, approximately 20,000 frames were acquired using simultaneous excitation with 560 nm and 647 nm lasers with an exposure time of 100 ms. Imager strand solutions of Cy3B and ATTO 647N were thoroughly mixed using a pipette before being introduced into the microfluidic device, with final concentrations of 0.01 nM and 0.001 nM, respectively.

For single-target biplane imaging of mitochondria, approximately 40,000 frames were acquired using 560 nm and 647 nm laser excitation with an exposure time of 100 ms. Biplane detection was implemented by axially shifting the second relay lens in the red detection path by ∼29 mm toward the camera, thereby generating two offset image planes on the sCMOS sensor. During biplane alignment, special care was taken to ensure that the DH phase mask remained aligned in the Fourier plane throughout the lens translation, as any displacement of the phase mask would introduce additional aberrations and compromise localization precision and accuracy.

For dual-target single-plane single-molecule imaging of mitochondria and the nuclear lamina, approximately 40,000 frames were acquired using 560 nm and 647 nm laser excitation with an exposure time of 100 ms per frame. The emitted fluorescence was separated into green and red spectral channels using a dichroic mirror. Both channels were projected onto separate regions of the same sCMOS sensor, ensuring simultaneous acquisition of the two color channels without axial plane offset. To confirm there is no axial plane offset we used fiducial beads and DH-PSFs to align the channels.

### Light-sheet versus epi-fluorescence performance evaluation

The performance of the LS illumination was quantitatively evaluated against conventional epi-fluorescence illumination in both the 560 nm and 647 nm spectral channels using fluorescently labeled lamin B1. Imaging was performed under identical detection settings, with exposure time of 50 ms and laser power of 150 mW with ND 2.6 at the laser output. For each excitation wavelength, 10 images were acquired sequentially with z lock (PFS on, Nikon) under LS and epi-illumination. The images acquired were then averaged and merged using ImageJ to get the comparison of epi- versus LS illumination. Intensity line profiles were generated across representative nuclear regions from the images. The data showed ∼3-3.5-fold and ∼2-2.5-fold improvements in the SBR when switching from epi- to LS illumination for the 560 nm and 647 nm laser, respectively (Supplementary Fig. 13).

Single molecule DNA-PAINT data of lamin B1 in U2OS cells was acquired in 3D using the DH-PSF with epi- or LS illumination to assess the performance of the SoLiD-3D platform in terms of localization precision (Supplementary Fig. 14). The background photons per pixel (px), detected signal photons per localization, and localization precision (σ_xy_ and σ_z_) were extracted using Easy-DHPSF.

Under 560 nm excitation, the range of median values from three replicates for LS/epi were 11–14/19-25 background photons per pixel, 1933–2940/1718-2271 signal photons per localization, 21–24/21–27 nm lateral localization precision, and 24–33/31–42 nm axial localization precision.

Under 647 nm excitation, the range of median values from three replicates for LS/epi were 31-37/42-58 background photons per pixel, 1915–3032/1882–3201 signal photons per localization, 20–27/22–29 nm lateral localization precision, and 31–41/33–45 nm axial localization precision.

### AutoDS3D based localization and computational analysis

Single-molecule localization of reconstructed DNA-PAINT data was performed using AutoDS3D^36,42^ neural network-based localization framework, with independent models trained separately for each PSF configuration (DH-3, DH-6, and Hummus-3) to account for their distinct characteristics and axial encoding behavior. Training datasets were generated from experimentally acquired fluorescent bead z-stacks, ensuring that the network learned realistic system-specific aberrations and optical signatures. Bead samples (TetraSpeck, 0.2 µm diameter, Invitrogen) were imaged under identical optical conditions, with axial scans acquired in 100 nm steps over ranges of −1.5 to +1.5 µm for DH-3 and Hummus-3, and −3 to +3 µm for DH-6. The experimentally measured PSF stacks were used to simulate emitter images by sampling random 3D emitter coordinates (x, y, z) within the calibrated axial range (∼10,000 images, 1-35 molecules, 13.1 × 13.1 µm² area). To ensure realistic training conditions, both signal and background noise were calibrated on experimentally measured single-molecule photon counts and noise statistics, thereby closely matching real acquisition conditions.

For experimental data processing, raw image sequences were first subjected to temporal background subtraction. Background subtraction could be implemented in either of two ways. In the first approach, a rolling average was computed and subtracted from each individual frame, providing a temporally smoothed estimate of the background, or images were processed in groups of 100 frames, where a pixel-wise minimum intensity projection was generated for each group and subsequently subtracted from the corresponding individual frames. Both approaches effectively removed slowly varying background contributions while preserving single-molecule emission signals. The resulting background-subtracted frames were processed using the corresponding AutoDS3D model selected according to the PSF used during acquisition. A detection threshold was empirically optimized for each dataset such that the resulting reconstructions achieved a Jaccard index in the range of 0.4–0.7, balancing precision and recall across varying emitter densities. Inference was performed and produced various localization outputs for each emitter including *x*, *y*, and *z* coordinates, frame number, and intensity. The resulting localization tables were exported as .CSV files for post-processing.

Green and red channel localization datasets were registered using affine transformation parameters obtained from calibration bead datasets to map the red channel onto the green channel, and further refined using cross-correlation optimization to achieve sub-diffraction alignment accuracy.

To correct sample drift, fiducial beads (T7280, TetraSpeck, 0.2 μm diameter; Invitrogen) that remained visible throughout the acquisition were used as a stable spatial reference. In the green channel, bead positions were directly tracked either using Easy-DHPSF^47^ or AutoDS3D^42^ and extracting the bead cluster in the x–y localization map, allowing only localizations corresponding to the same bead to be retained. The selected bead localizations were then temporally filtered using a sliding frame window to suppress outlier localizations and reduce frame-to-frame localization noise. The filtered x, y, and z coordinates were subsequently fit with cubic smoothing splines to estimate a continuous three-dimensional drift trajectory over the full acquisition. The smoothing parameters were chosen independently for each spatial dimension to capture slow mechanical and thermal drift while suppressing stochastic localization fluctuations. The drift at each frame was calculated as the deviation of the smoothed bead trajectory from its initial position and was then subtracted frame-by-frame from the corresponding single-molecule localizations in x, y, and z. Custom-written MATLAB codes were used for all registration and drift correction operations.

The final merged and drift corrected localization datasets were imported into the Vutara SRX software (Bruker) for visualization and rendering, where point splat representations were generated with Gaussian rendering, molecule size as 50 nm, opacity of 0.7-0.8 and axial color-coding to reflect z-position. The mitochondria reconstructions were filtered to remove spurious localizations based on the method of mean distance with a distance threshold of 13-14 nm and 20 nearest neighbors. and The Fourier ring correlation (FRC) was calculated in Vutara SRX to estimate the achieved 3D spatial resolution of the single-molecule localization microscopy datasets. By splitting the localized data into independent subsets, the software plots correlation curve against spatial frequency across the lateral (xy) and axial (xz and yz) planes using an 8 nm pixel size binning, determining the resolution at the point where this curve drops below the a fixed (1/7) threshold.

### CRLB analysis of depth-dependent localization performance

The theoretical localization precision of the engineered point spread function (PSF) was evaluated using Cramér–Rao lower bound (CRLB) analysis. The phase mask used in the optical model was generated from the calibration z-stack of experimental data using the VIPR framework^37^. The resulting phase mask array was saved in a .mat file and subsequently imported into the simulation pipeline. Single emitters were simulated over an axial range of −1.5 to +1.5 µm with 50-nm spacing, while the lateral position was fixed at the optical axis (x = 0, y = 0). Each emitter was assigned a photon budget of 6,000 detected photons, and the corresponding PSFs at each axial position were generated using a forward imaging model incorporating the optimized phase mask. To quantify the sensitivity of the PSF to emitter displacement, required for Fisher information analysis, the emitter position was independently perturbed by 1 nm along the x, y, and z directions, generating additional PSFs at (x+dx, y, z), (x, y+dy, z), and (x, y, z+dz), where dx = dy = dz = 1 nm. Numerical partial derivatives of the expected photon distribution with respect to each spatial coordinate were estimated using first-order finite differences between the perturbed and unperturbed PSFs, thereby quantifying the response of the image intensity distribution to sub-nanometer emitter displacements. The Fisher information matrix (FIM) was computed independently at each axial position under the assumption of Poisson-distributed photon detection with a spatially mean background of 10 photons per pixel. Individual matrix elements were obtained by summing pixel-wise products of the corresponding spatial derivatives, normalized by the expected photon counts at each detector pixel. The inverse of the FIM provided the lower bound on the covariance of the estimated emitter position, and the square root of its diagonal elements was reported as the theoretical localization precision in x, y, and z. The resulting localization precisions were converted to nanometers and plotted as a function of axial position to quantify depth-dependent localization performance. Representative PSFs at selected axial positions were additionally visualized to examine the evolution of PSF morphology across the imaging volume. Performance under reduced photon counts (3,000 signal photons per localization) and elevated background levels (30 background photons per pixel) is additionally characterized (Supplementary Fig. 15).

### Jaccard index and localization error analysis

To quantitatively evaluate localization performance across different emitter densities, we used a simulation-based benchmarking pipeline with experimentally measured PSF models and a known ground-truth emitter generator^36^. For each condition, the number of emitters was systematically varied across a predefined range (2–180 emitters per field of view), corresponding to a density range of ∼0 to 1.1 µm⁻², where density was computed based on the effective imaging area of 13.1

× 13.1 µm². For each density condition, 100 independent Monte-Carlo realizations were generated to ensure statistical robustness. For each realization, ground-truth emitter coordinates (*x*, *y*, *z*) were sampled using a stochastic emitter generator constrained by the experimental PSF model. Corresponding simulated images were generated, incorporating experimentally calibrated photon statistics, background levels, and camera noise parameters, which are defined and maintained during DeepSTORM3D model training. These simulated images were then passed through the trained DeepSTORM3D inference network to obtain predicted emitter coordinates and associated confidence values.

Performance was quantified using a custom matching-based evaluation routine that computed the Jaccard index and RMSEs in the lateral (*xy*) and axial (*z*) dimensions. The Jaccard index was defined as the ratio between true positive localizations and the union of detected and ground-truth emitters, J=TP/(TP+FP+FN), where TP is true positive, FP is false positive, and FN is false negative), where true positives were determined using a nearest-neighbor assignment between predicted and ground-truth coordinates with a maximum matching tolerance of 0.1 µm. Specifically, a predicted localization was considered a true positive if its Euclidean distance from a ground-truth emitter fell within this threshold, unmatched predictions were counted as false positives, and unmatched ground-truth emitters as false negatives. This formulation provided a robust measure of detection accuracy under increasing emitter overlap conditions. Both the Jaccard index and RMSE are strongly dependent on photon budget and background conditions. The results reported here correspond to a high-photon regime representative of optimal imaging conditions. Performance under reduced photon counts and elevated background levels is additionally characterized (Supplementary Fig. 16).

Localization precision was further quantified using RMSE metrics computed separately for lateral and axial coordinates. For matched emitter pairs, the RMSE in the *xy*-plane was computed as

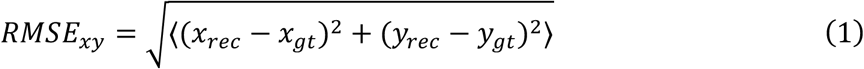

where, RMSE_xy_ is root mean square error in the lateral (*xy*) plane, x_rec_ is reconstructed *x*-coordinate of the emitter, *y*_rec_ is reconstructed *y*-coordinate of the emitter, *x*_gt-_ is the ground-truth *x*-coordinate of the emitter and *y*_gt-_ is the ground-truth *y*-coordinate of the emitter and the axial error was computed as

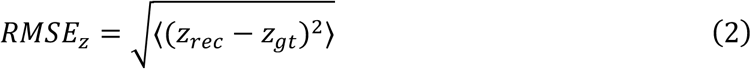

where RMSE_z_ is root mean square error along the axial (*z*) direction, *z*_rec_ is reconstructed axial position of the emitter, and *z*_gt_ is ground-truth axial position of the emitter.

For each emitter density, the Jaccard index and RMSE values were computed independently for all 100 runs, and the final reported metrics correspond to the mean and standard deviation across realizations. In cases where no valid localization was returned by the network, the Jaccard index was set to zero and RMSE values were excluded from averaging using NaN-based statistical handling.

### PSF footprint analysis and emitter density estimation

We further quantitatively compared the packing density achievable with Hummus-3 versus the DH-3 and DH-6. To estimate the maximum resolvable emitter density, we used experimentally calibrated PSFs for all three PSFs (Supplementary Fig. 17). Experimental PSFs were obtained using fluorescent beads illuminated with epi-illumination. For each PSF design, calibration stacks were recorded over an axial range of 3 µm with an axial step size of 100 nm. For packing analysis, each PSF image was normalized and converted into a binary image using a fixed intensity threshold of 0.7 % which was applied identically to all PSF designs to define the effective spatial footprint. Pixels with intensities above the threshold were assigned a value of 1, while all remaining pixels were assigned a value of 0. The resulting binary mask represented the minimum detector area occupied by a single emitter. To estimate the maximum number of non-overlapping emitters that could be simultaneously accommodated within a single camera frame, each binary PSF was computationally packed into a fixed image region of 121 × 121 pixels (pixel size = 108 nm, corresponding field of view = 13.1 µm × 13.1 µm) using a custom MATLAB algorithm. During each iteration, emitter positions were randomly generated within the available image area. A new PSF was accepted only if no pixel overlap occurred with previously placed masks. The procedure was repeated iteratively until no additional PSFs could be placed after that the process was terminated. The maximum number of pixels that can be occupied was kept as constant for all three phase masks (∼8,000). The total number of successfully placed masks was recorded as the maximum geometric packing density for that PSF design.

To further evaluate how the effective PSF footprint varied as a function of axial position, binary PSFs were generated independently for each calibrated z-plane across the full measurement range (0-3 µm). The total number of foreground pixels occupied by each PSF at axial position *z* was calculated as

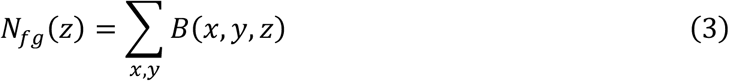

where *B*(*x*, *y*, *z*) denotes the binarized PSF mask at axial position. Foreground pixel counts were then compared across all PSF designs to quantify changes in spatial footprint throughout the calibrated imaging volume. All image processing, binarization, logical mask operations, and iterative packing simulations were performed using custom scripts implemented in MATLAB.

## Author contributions

P.J. and A.-K.G. conceived the idea. P.J. constructed the imaging platform and developed the data analysis pipeline. P.J, N.S. and S.C. fabricated microfluidic chips. P.J., S.C., N.S., and Y.N. prepared samples for DNA-PAINT imaging. P.J., Y.N., and D.X. modified DeepSTORM3D and Auto-DeepSTORM3D code for analyzing Hummus-3. P.J., S.C., and N.S. wrote and modified codes for camera triggering, registration, and stitching. D.X., R.O., and Y.S. designed and fabricated the Hummus-3 phase mask. A.-K.G. and Y.S. provided supervision and intellectual discussions on the research. P.J. and A.-K.G. wrote the manuscript with input from all authors. All authors discussed the data and contributed to editing the manuscript.

## Data and code availability statement

Source data that can be used for generating the figures and graphs in this work are provided with this paper and on GitHub [https://github.com/Gustavsson-Lab/SoLiD-3D].

Calibration and fitting analysis of DH-PSF images for epi- versus LS comparisons was performed using a modified version of the open-source Easy-DHPSF software^44,47^

[https://sourceforge.net/projects/easy-dhpsf/]. The Hummus-3 phase masks were designed using DeepSTORM3D^36^ [https://github.com/EliasNehme/DeepSTORM3D]. All single-molecule data analysis except for the epi- versus LS comparison, as well as Jaccard index analysis, was performed using AutoDS3D^42^ [https://github.com/dafeixiao/AutoDS3D]. The custom-written codes for red-to-green channel transformation, drift correction, channel cross-correlation, sCMOS camera control via Micro-Manager^41^, galvanometric mirror control, and image processing, binarization, logical mask operations, and iterative packing simulations for the PSF footprint and density evaluation are available on Github [https://github.com/Gustavsson-Lab/SoLiD-3D].

## Acknowledgements

We thank Tyler Nelson for help in 3D Blender, Margareth Freire for help with sample labeling, and Parth Joshi for help with initial data acquisitions. Sample chambers in this work were fabricated, imaged, and profiled using resources and equipment available through the Shared Equipment Authority at Rice University.

