## Supplementary Information for "High-speed volumetric single-molecule imaging using dual-wavelength light sheets and PSF-engineered enhanced biplane detection"

**Table of contents**

**Figure S1: Detailed schematic and rendering of the optical setup.**

**Figure S2: Light-sheet (LS) thicknesses**

**Figure S3: Light-sheet (LS) illumination with and without dithering**

**Figure S4: Confocal parameters of the light sheets (LSs)**

**Figure S5: Light-sheet (LS) focusing characterization**

**Figure S6: Axial calibration of the DH-3 PSF and 4f lens displacement**

**Figure S7: Fourier ring correlation (FRC) analysis corresponding to mitochondrial reconstruction data in Figure 2**

**Figure S8: Fourier ring correlation (FRC) analysis corresponding to mitochondrial reconstruction data in Figure 3**

**Figure S9: Fourier ring correlation (FRC) analysis corresponding to nuclear lamina and mitochondrial reconstruction data in Figure 4**

**Figure S10: Comparative PSF footprint and foreground pixel occupancy**

**Figure S11: Fourier ring correlation (FRC) analysis corresponding to mitochondrial reconstruction data in Figure 7**

**Figure S12: Galvanometric mirror calibration for dual-wavelength light-sheet (LS) translation**

**Figure S13: Enhanced optical sectioning improves the signal-to-background ratio**

**Figure S14: Enhanced optical sectioning improves the localization precision**

**Figure S15: PSF design, characterization, and precision benchmarks under low-photon conditions**

**Figure S16: Comparative performance of engineered and learned PSFs under low-photon conditions**

**Figure S17: Experimental images of the Hummus-3, and Double-Helix (DH)-3 and DH-6 PSFs across different axial ranges**

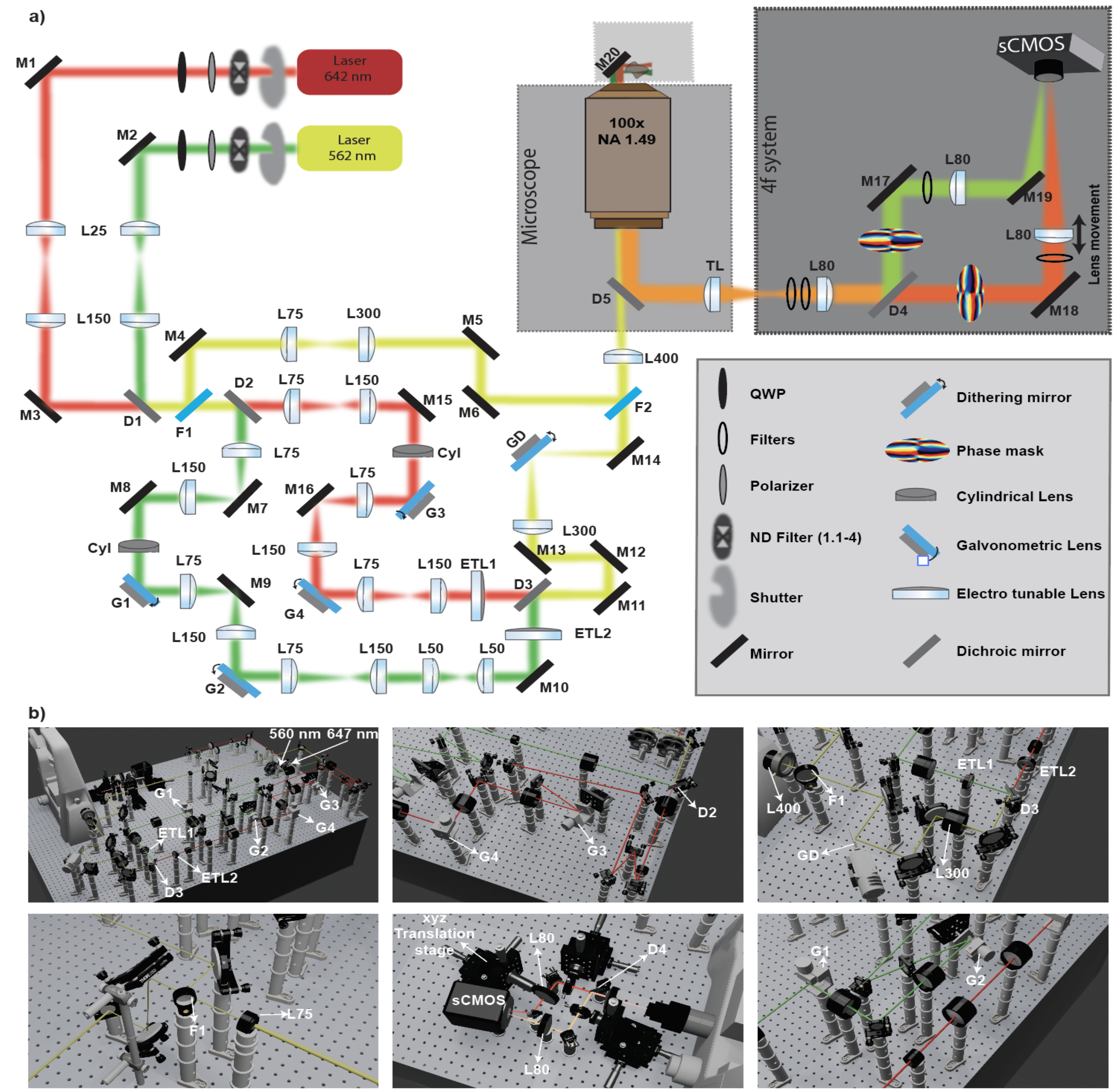


**Figure S1: Detailed schematic and rendering of the optical setup.** (a) Detailed schematic of the optical setup. The schematic is not to scale. (b) To-scale rendering showing different views of the setup.

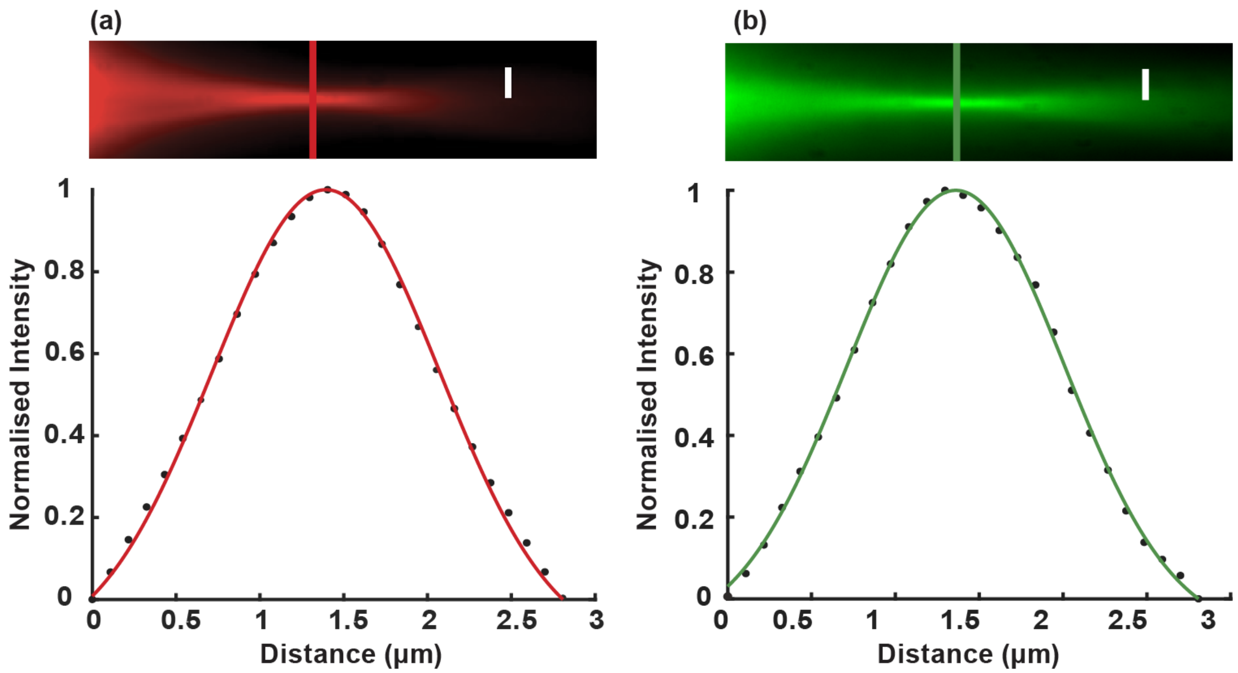


**Figure S2: Light-sheet (LS) thicknesses.** (a-b) The narrow ends of the LSs imaged in a fluorescent solution (Cy3B for 560 nm, ATTO 647N for 647 nm). The graphs show the corresponding line profiles with Gaussian fits overlaid. The LS thicknesses ($1/e^2$ diameter) were measured as (a) 2.3 µm for 647 nm and (b) 2.4 µm for 560 nm, corresponding to beam waist radii of approximately 1.2 µm. Scale bars: 5 µm.

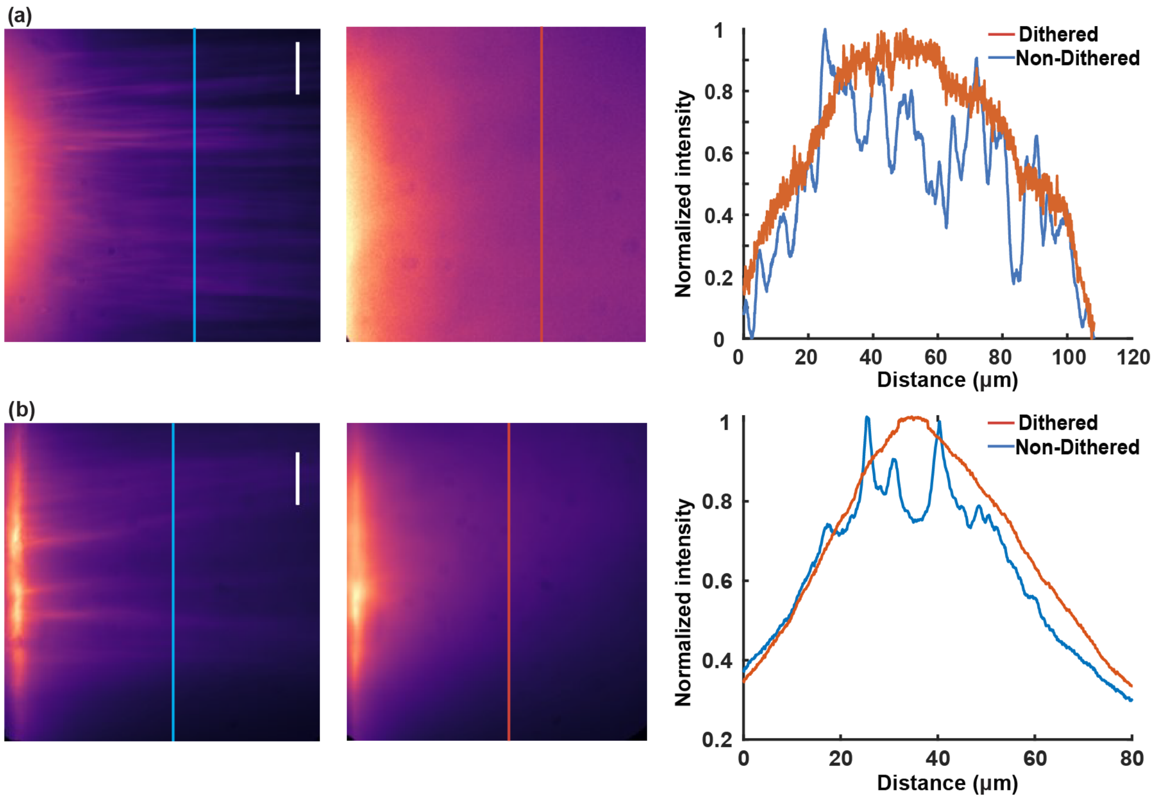


**Figure S3: Light-sheet (LS) illumination with and without dithering.** (a–b) The left column shows striping artifacts observed with non-dithered LS illumination in fluorescent solutions containing (a) Cy3B for 560 nm, and (b) ATTO 647N for 647 nm. The middle column demonstrates homogeneous illumination achieved by dithering the LS within the illumination plane using a galvanometric dithering mirror operated at 100 Hz. Scale bars: 10 µm. The graphs show line scans at the indicated lines in the images. The measured dithered LSs widths ($1/e^2$ diameter) were (a) ~56 µm at 560 nm and (b) ~48 µm at 647 nm.

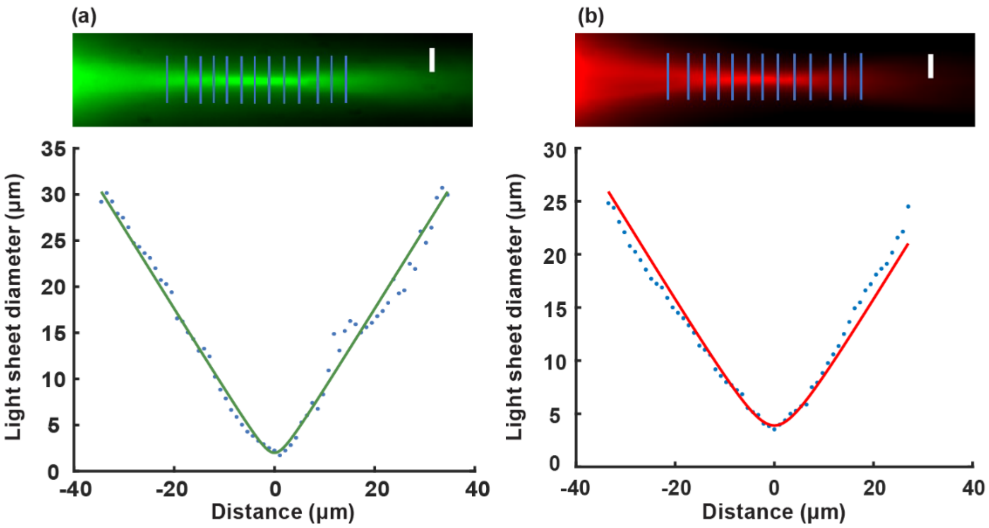


**Figure S4: Confocal parameters of the light sheets (LSs).** The confocal parameters of the LS were measured by imaging the narrow ends of the LSs in fluorescent solutions containing (a) Cy3B for 560 nm, and (b) ATTO 647N for 647 nm. Gaussian fits to the axial beam profiles at different positions along the propagation direction were used to determine the ranges over which the LSs maintained a thickness within $\sqrt{2}$ of their thickness at focus. The resulting confocal parameters were (a) 15.3 µm for 560 nm and (b) 13.2 µm for 647 nm, representing the effective sectioning range along the direction of beam propagation. Scale bars: 5 µm.

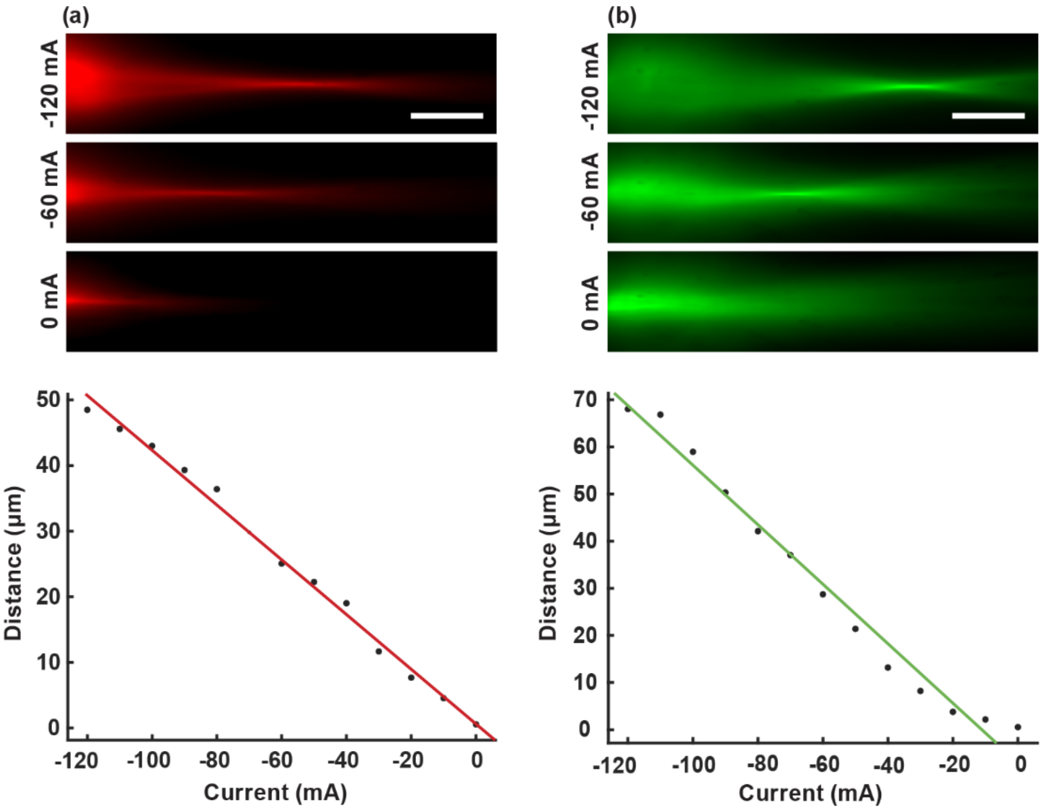


**Figure S5: Light-sheet (LS) focusing characterization.** (a–b) Representative images of the (a) red (647 nm) and (b) green (560 nm) LSs in fluorescent solution containing Cy3B for 560 nm and ATTO 647N for 647 nm, with corresponding graphs showing the displacement of the LS foci as a function of current applied to the tunable lenses. The lens currents were varied from 0 mA to –120 mA in 10 mA increments. Scale bars: 10 µm.

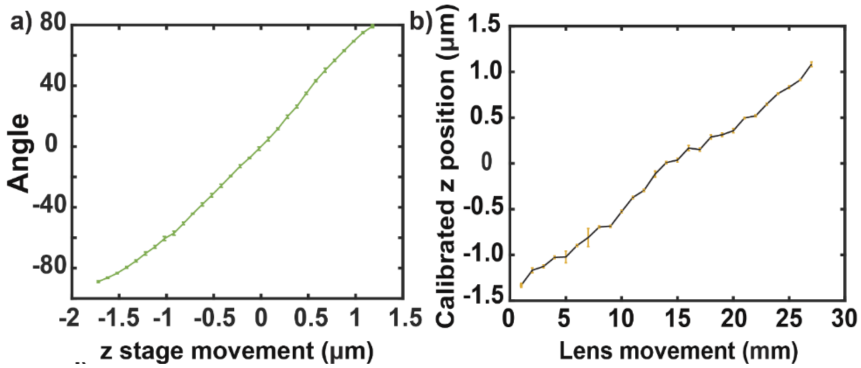


**Figure S6: Axial calibration of the DH-3 PSF and 4f lens displacement**. (a) Graph showing calibration of the DH-3 engineered PSF rotation angle as a function of emitter axial position (z), enabling angle-encoded axial localization of single molecules. (b) The second lens in the red 4f detection path is mounted on a translational stage, allowing controlled axial displacement toward and away from the camera. The graph shows the axial position (z) calibrated as a function of lens displacement, with the angle–z relationship serving as the reference to define the axial response to lens motion. A displacement of 1 mm of the lens corresponds to an axial shift of approximately 100 nm in the sample plane. The plots show the mean ± standard deviation of three replicated measurements.

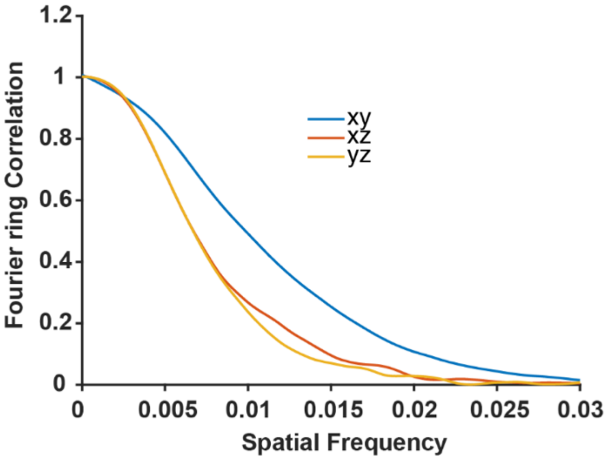


**Figure S7. Fourier ring correlation (FRC) analysis corresponding to mitochondrial reconstruction data in Figure 2.** FRC curves plotted as a function of spatial frequency to assess the resolution of the 3D dataset.

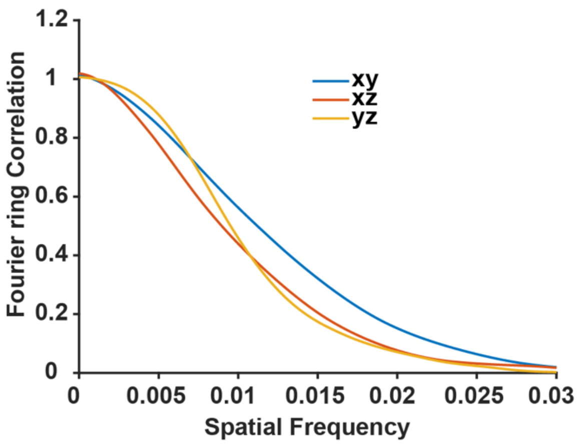


**Figure S8. Fourier ring correlation (FRC) analysis corresponding to mitochondrial reconstruction data in Figure 3.** FRC curves plotted as a function of spatial frequency to assess the resolution of the 3D dataset.

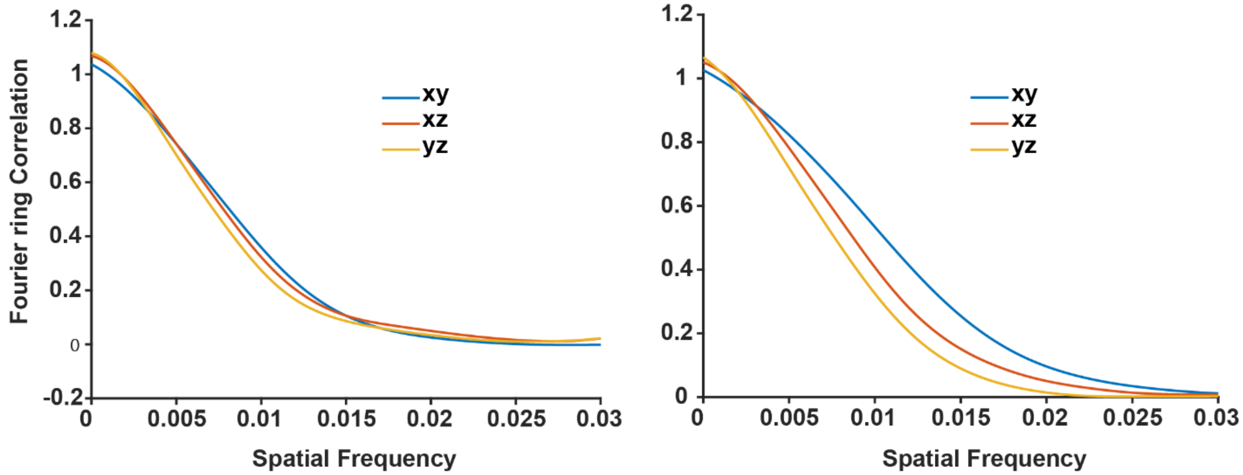


**Figure S9. Fourier ring correlation (FRC) analysis corresponding to nuclear lamina and mitochondrial reconstruction data in Figure 4.** FRC curves plotted as a function of spatial frequency to assess the resolution of the 3D datasets, showing the nuclear lamina data (left) and mitochondria data (right).

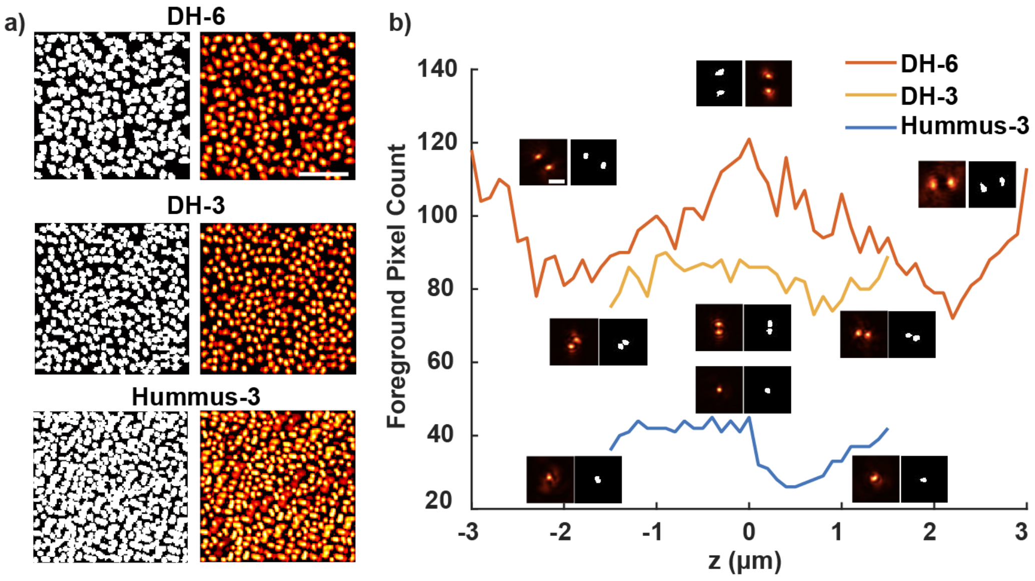


**Figure S10: Comparative PSF footprint and foreground pixel occupancy.** (a) Representative simulated widefield frames (right) and their corresponding binary segmentation masks (left) for the DH-6, DH-3, and Hummus-3 PSFs at a high emitter density. The masks illustrate the spatial extent and degree of overlap for each PSF design under identical imaging conditions. Scale bar: 5 µm. (b) Quantitative analysis of the PSF footprint, expressed as the foreground pixel count (the number of pixels above a defined intensity threshold) as a function of the axial position (z). The Hummus-3 PSF maintains a consistently smaller footprint across its entire 3 µm axial range compared to the DH-3 and DH-6 PSF designs. Insets show individual PSF snapshots alongside their binary masks at specific axial planes, highlighting the structural compactness of the learned Hummus-3 design. This reduced spatial occupancy directly correlates with a lower probability of emitter signal overlap, facilitating more accurate multi-emitter recovery in dense 3D SMLM samples.

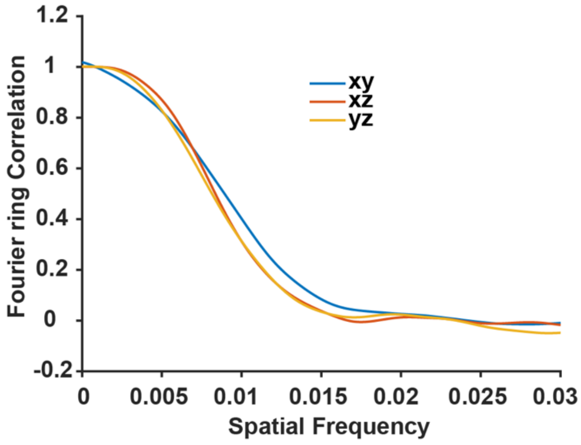


**Figure S11. Fourier ring correlation (FRC) analysis corresponding to mitochondrial reconstruction data in Figure 7.** FRC curves plotted as a function of spatial frequency to assess the resolution of the 3D dataset.

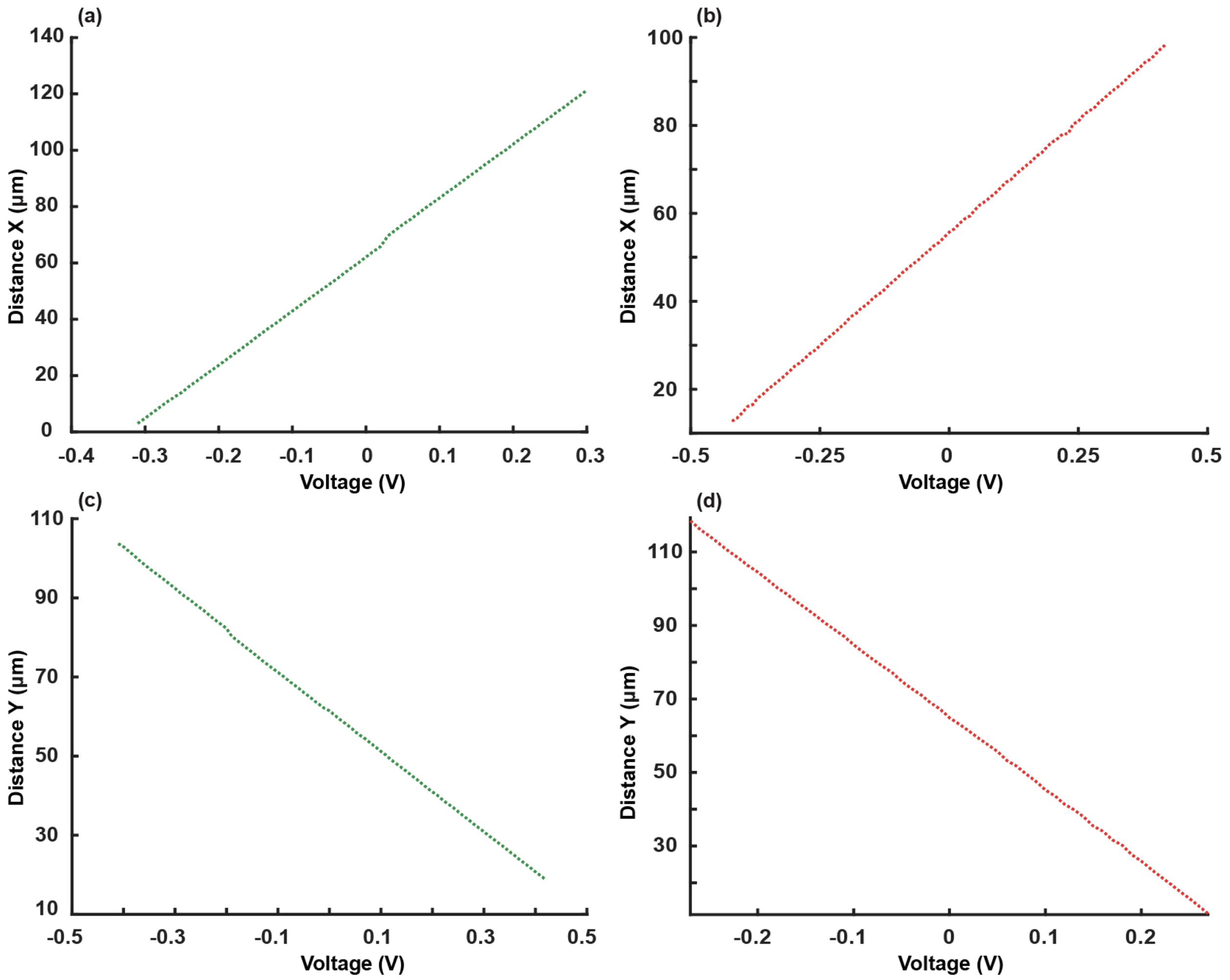


**Figure S12: Galvanometric mirror calibration for dual-wavelength light-sheet (LS) translation.** (a–b) Graphs showing the linear displacement along the x-axis as a function of applied voltage for the galvanometric X mirror, measured at (a) 560 nm and (b) 647 nm excitation wavelengths. (c–d) Graphs showing the linear displacement along the y-axis as a function of applied voltage for the galvanometric Y mirror, measured at (c) 560 nm and (d) 647 nm excitation wavelengths. The galvanometric mirrors translate the LSs in a highly linear fashion across both channels. For a voltage increment of 0.01 V, the LSs undergo a translation of approximately 1.07 µm (corresponding to 9.91 pixels based on a calibrated pixel size of 108 nm).

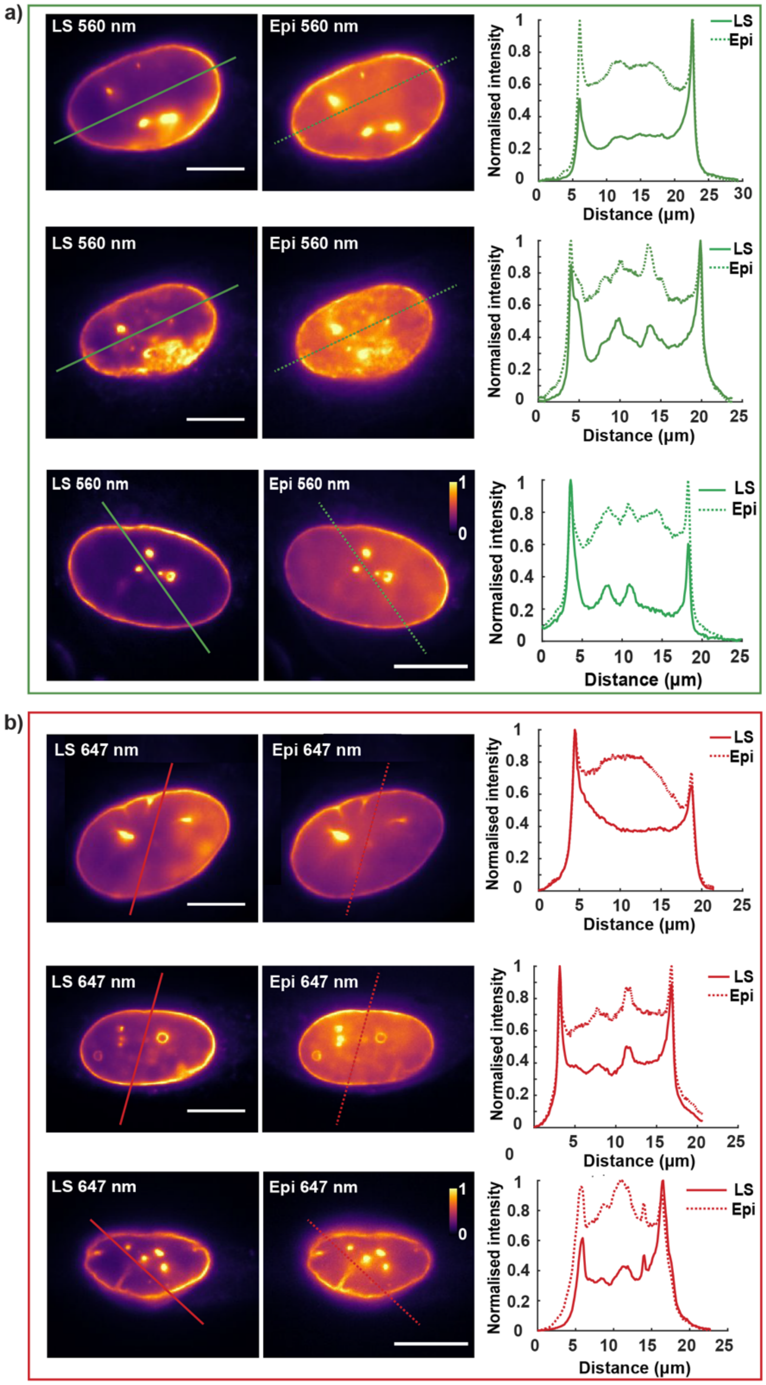


**Figure S13: Enhanced optical sectioning improves the signal-to-background ratio.** Diffraction-limited images of lamin B1 in U2OS cells excited with epi- or light sheet (LS) illumination when illuminated with (a) 560 nm and (b) 647 nm light. Graphs show line scans demonstrating the contrast improvement when using LS compared to epi-illumination. Scale bars 5 µm. The colorbars show intensity normalized independently for each image. Diffraction-limited imaging was repeated for $n = 3$ cells , demonstrating consistent improvements in signal-to-background ratio.

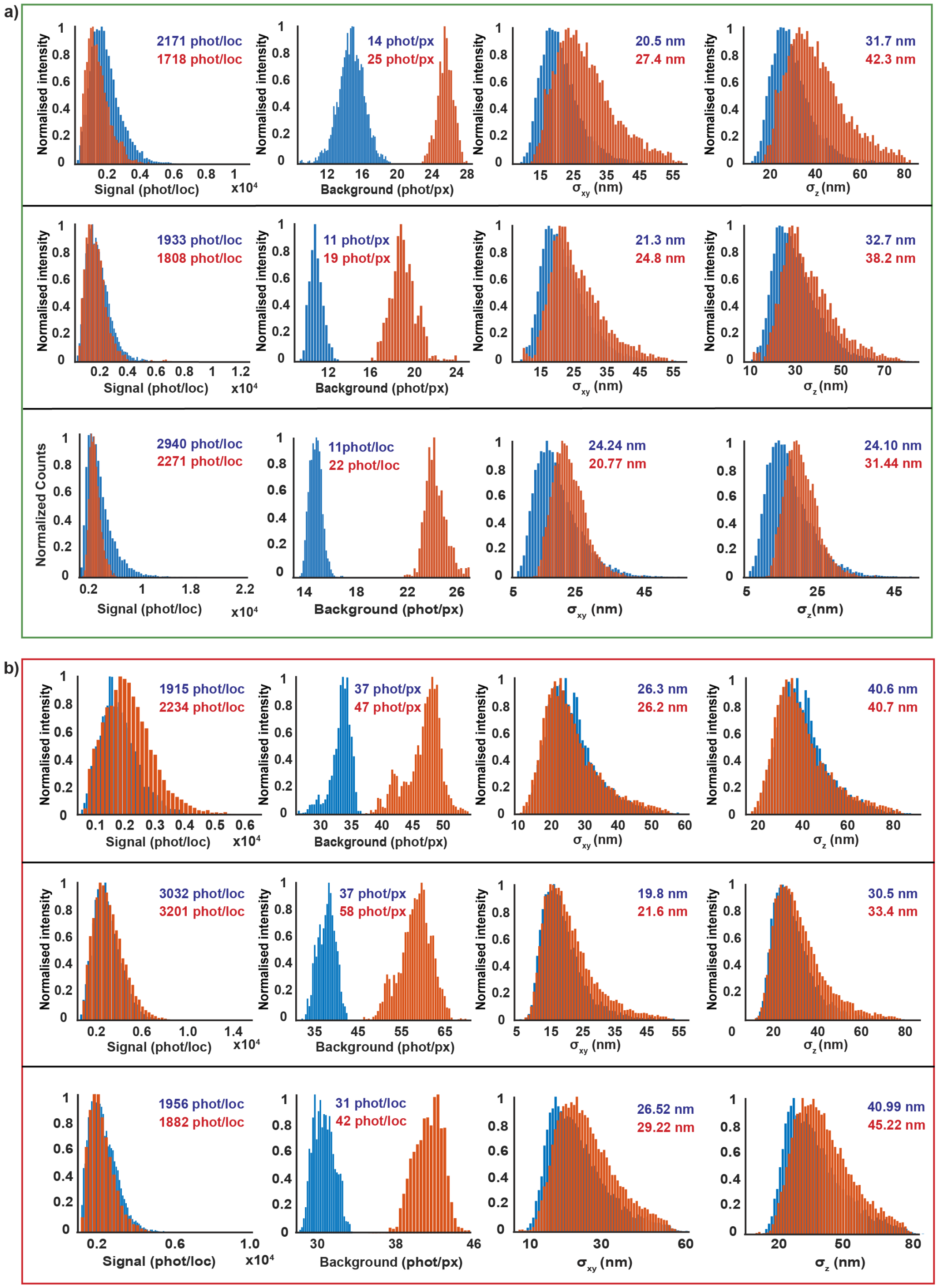


**Figure S14: Enhanced optical sectioning improves the localization precision.** Histograms demonstrating comparable signal photon levels but decreased fluorescence background levels leading to improved localization precisions in *xy* and *z* for LS compared to epi-illumination for 3D single-molecule super-resolution imaging of lamin B1 when using (a) 560 nm and (b) 647 nm light. The experiments were repeated for $n = 3$ cells, demonstrating consistent results.

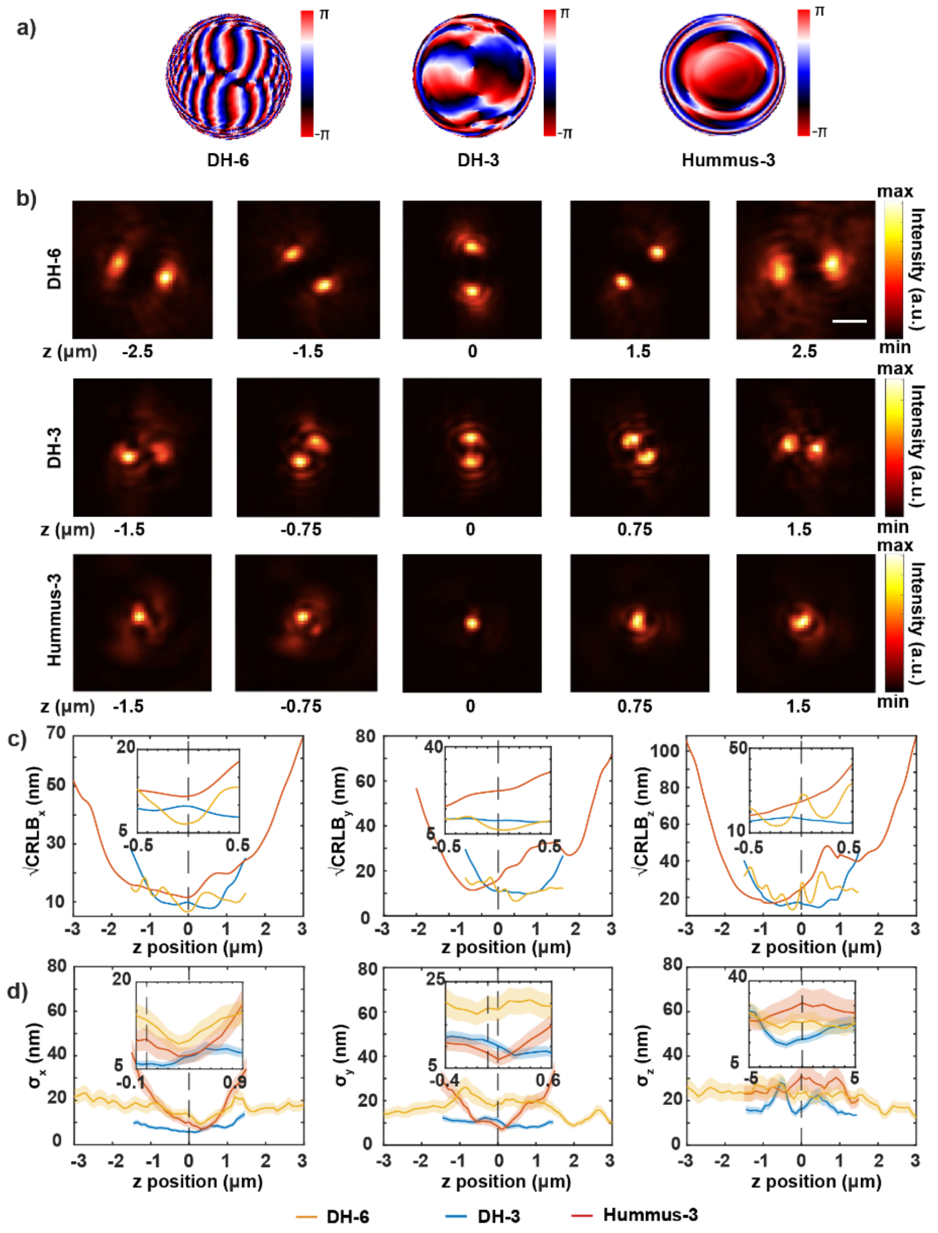


**Figure S15: PSF design, characterization, and precision benchmarks under low-photon conditions.** (a) Three phase masks were used and compared in this study: a double-helix (DH) phase mask designed for a 6 μm axial range (DH-6), a DH phase mask designed for a 3 μm axial range (DH-3), and a newly optimized Hummus phase mask designed for a 3 μm axial range (Hummus-3). The DH phase patterns were generated from experimentally obtained z-scans and implemented using the VIPR framework, whereas the Hummus-3 phase mask was designed using DeepSTORM3D. (b) Simulated PSFs for the DH-3, Hummus-3, and DH-6 phase masks, illustrating the relative PSF footprints and axial encoding characteristics. Scale bar: 2 μm. (c) Numerically calculated localization precision in the x, y, and z dimensions, defined as the square root of the Cramér–Rao lower bound ($CRLB^{1/2}$), computed from the simulated PSFs. Simulations used 3,000 signal photons and a mean background of 30 photons per pixel. (d) Experimental localization precision obtained from repetitive localization of fluorescent beads imaged with the three phase masks. All experimental measurements were acquired under the same conditions. Shaded areas represent the standard deviation of 100 measurements for each z.

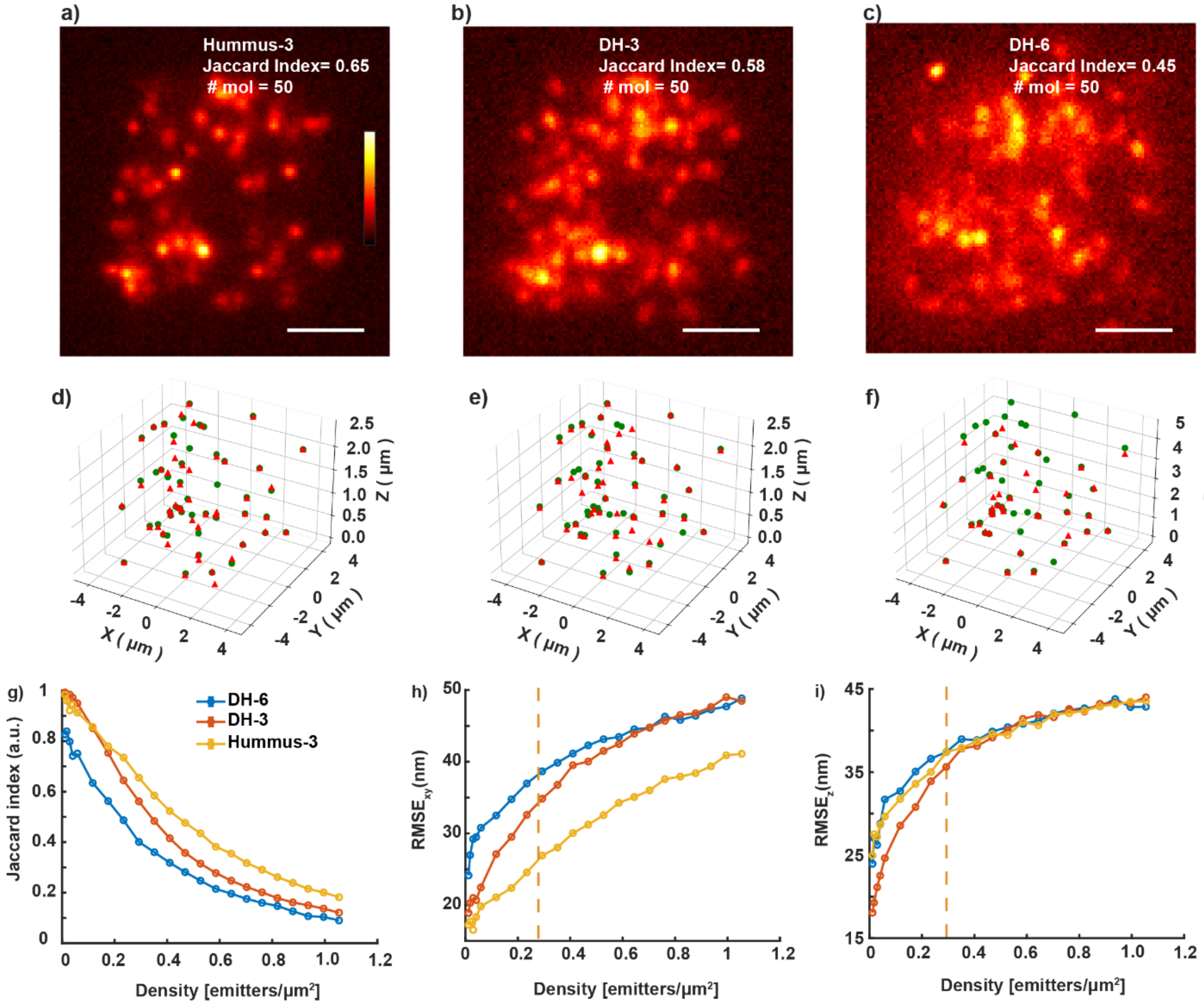


**Figure S16: Comparative performance of engineered and learned PSFs under low-photon conditions.** (a-c) Representative simulated images of 50 recognized molecules using the DH-6, DH-3, and Hummus-3 PSFs. The resulting Jaccard index is indicated for each PSF at an emitter density of 0.3 emitters/μm². Scale bars: 2 μm. (d-f) 3D scatter plots comparing ground truth (GT, green circles) with reconstructed (Rec, red triangles) molecule positions for the cases shown in (a-c), for DH-6 (J = 0.45, 62 detections, TP = 33, FP = 29, and FN = 18), for DH-3 ( J = 0.58, detections, TP = 42, FP = 15, and FN = 8), and for Hummus-3 (J = 0.65, 68 detections, TP = 46, FP = 22, and FN = 5). (g-i) Quantitative metrics as a function of emitter density (emitters/μm²): (g) Jaccard index (detection accuracy), and root mean square error (RMSE) in the (h) lateral plane and (c) axial dimension. The vertical dashed orange line indicates a reference density of 0.3 emitters/μm². Abbreviations: J: Jaccard index, TP: true positives, FP: false positives, FN: false negatives.

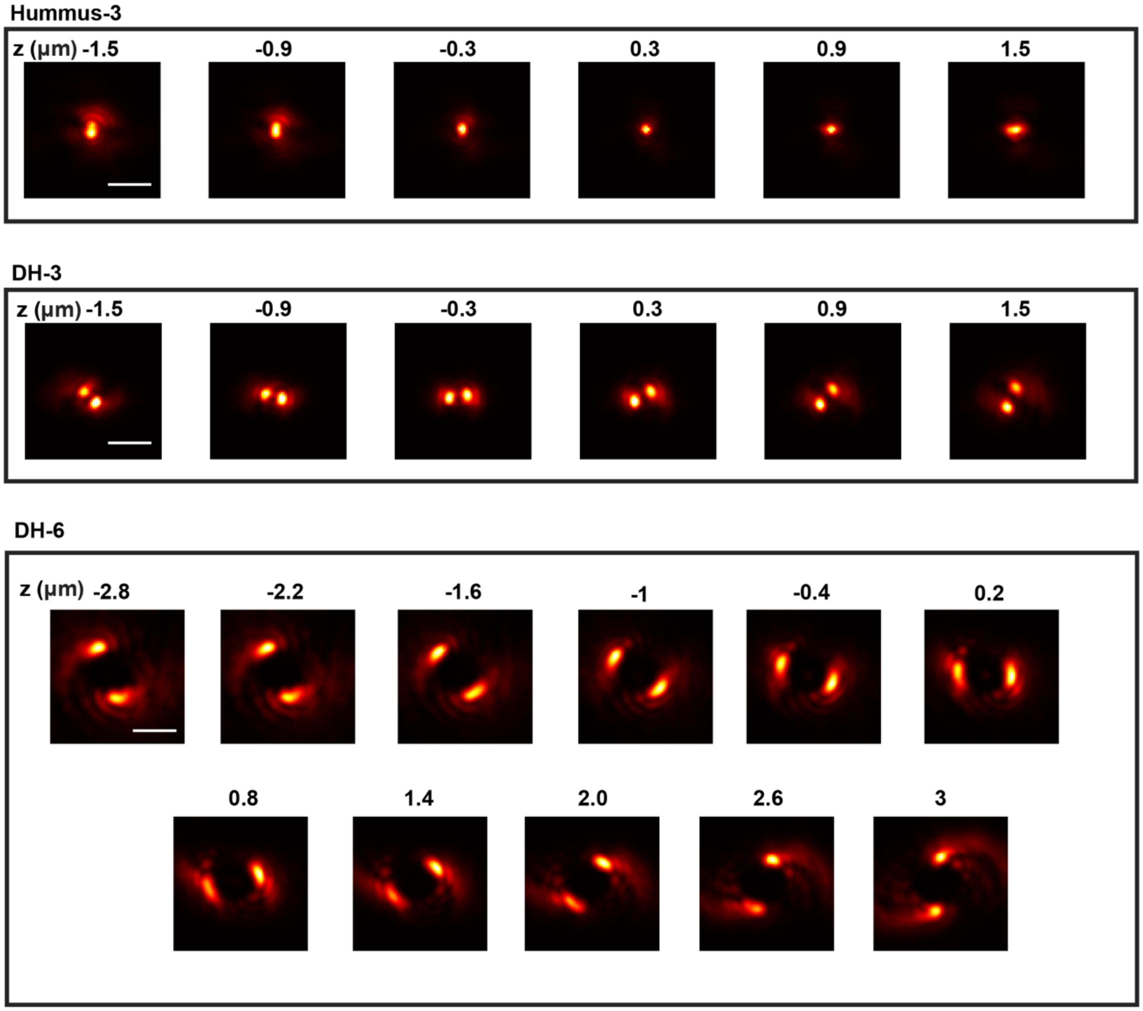


**Figure S17: Experimental images of the Hummus-3, and Double-Helix (DH)-3 and DH-6 PSFs across different axial ranges.** Experimental images of (a) the Hummus-3 PSF, (b) the DH-3 PSF, and (c) the DH-6 PSF across the indicated axial ranges. Scale bars: 3 μm.
